# Long-term stress and trait anxiety alter brain network balance in dynamic decisions during working memory

**DOI:** 10.1101/2021.05.07.442883

**Authors:** Liangying Liu, Jianhui Wu, Haiyang Geng, Chao Liu, Yuejia Luo, Jing Luo, Shaozheng Qin

## Abstract

Long-term stress has a profound impact on the human brain and cognition, and trait anxiety influences stress-induced adaptive and maladaptive effects. However, the neurocognitive mechanisms underlying long-term stress and trait anxiety interactions remain elusive. Here we investigated how long-term stress and trait anxiety interact to affect dynamic decisions during working-memory (WM) by altering functional brain network balance. In comparison to controls, male participants under long-term stress experienced higher psychological distress and exhibited faster evidence accumulation but had a lower decision-threshold during WM. This corresponded with hyper-activation in the anterior insula, less WM-related deactivation in the default-mode network, and stronger default-mode network decoupling with the frontoparietal network. Critically, high trait anxiety under long-term stress led to slower evidence accumulation through higher WM-related frontoparietal activity, and increased decoupling between the default-mode and frontoparietal networks. Our findings provide neurocognitive evidence for long-term stress and trait anxiety interactions on executive functions with (mal)adaptive changes.

## Introduction

Now, more than ever, increasing exposure to psychosocial stress has become an unavoidable part of our contemporary society as the pace of life rapidly accelerates. Exposure to sustained stress can have a profound impact on the brain and cognition through activation of stress-sensitive neuromodulatory systems and the release of stress hormones^1, 2^. The adverse effects of long-term stress on higher-order cognitive functions are widely documented^2, 3^. Yet, long-term stress has also been associated with no impairment or even enhanced cognitive functions^4^. Thus, long-term stress can lead to either beneficial forms of learning that promote adaptation or detrimental effects that presage maladaptation, likely depending on the level of stress resilience or vulnerability of each individual^5, 6^. Trait anxiety, a stable disposition to interpret a wide range of environmental events in a negative way, has been recognized as a vulnerable factor^7, 8^, which could account for the seemingly paradoxical effects of stress. Neurocognitive models of human anxiety suggest that high trait-anxious individuals tend to make additional effort to prevent shortfalls in performance effectiveness (i.e., accuracy) with deficits becoming evidence in processing efficiency (i.e., reaction times, RTs) on tasks involving executive function and deliberate dynamic decision processing such as WM ^9–11^. However, how trait anxiety and long-term stress interact to affect dynamic decisions during WM remains unclear. Considering both long-term stress and trait anxiety provides a better understanding of the profound effects of long-term stress on the brain and cognition than either in isolation.

Recent advances in computational modeling of trial-by-trial decision making enable us to identify latent dynamic computations in various cognitive domains including WM^12, 13^. Research in sequential-sampling theory posits that WM (i.e., the N-back task), analogous to speeded decision-making, can be modeled as an evidence accumulation process during which effective information (namely evidence) extracted from a stream of inherently noisy observations are rapidly accumulated until sufficient evidence reaches the threshold to make a decision and the choice response is then executed^14^. Drift diffusion model (DDM), in particular, is used to decompose participant’s choice responses in a given task into latent decision-making dynamics modulated by several free parameters. Of these parameters, the speed of evidence accumulation refers to ‘drift rate’ reflecting the ability to extract effective information from perceived inputs^15^, and ‘decision threshold’ represents the amount of accumulated evidence to reach a decision. The frontal-parietal network (FPN) regions, particularly the dorsolateral prefrontal cortex (dlPFC) in the middle frontal gyrus and the inferior parietal sulcus (IPS), are responsible for evidence accumulation^16^. Single-cell recordings in non-human primates have also established a link between latent evidence accumulation and neural firing rates in the dlPFC and the IPS^17–19^. These FPN regions are also the major targets of stress hormones such as glucocorticoids and catecholamines through which neuronal excitability and network connectivity in the PFC are affected^20, 21^. As a vulnerable phenotype of stress-related mental illness, high trait anxiety has been linked to deficient processing efficiency anchored onto the FPN critical for executive function^22^. Yet, how long-term stress and trait anxiety interact to affect the FPN in the dynamic decision process remains open.

The engagement of FPN regions during WM is usually accompanied by disengagement of core regions of the default mode network (DMN), especially the posterior cingulate cortex and medial prefrontal cortex. These DMN regions has been implicated in mind-wandering and allocation of resources to internal thoughts^23^. Greater disengagement of the DMN regions has been linked to better WM performance^24^, suggesting its role in suppression of irrelevant thoughts to support externally goal-directed tasks. Less DMN deactivation has been observed under acute stress and anxious individuals^25^, implying deficient reallocation of resources from internal thoughts to external tasks. Moreover, the salience network (SN) has been implicated in triggering a shift of neurocognitive resources to prioritize affective processing over deliberate executive functions including WM^26^. Indeed, hyper-activation in the anterior insula and dorsal anterior cingulate cortex (dACC), core nodes of the SN, have been reported by many studies in anxious individuals^23^. Although reduced disengagement of the DMN and hyper-activation in the SN has been observed under acute stress^27^, whether or not sustained exposure to stress leads to a similar effect on the DMN and SN remains unclear.

Beyond local activation, human WM relies on nuanced functional coordination among large-scale brain networks of the FPN, SN and DMN to support constantly maintaining and updating of relevant information according to ever-changing cognitive/environmental demands. In particular, functional antagonism or decoupling between FPN and DMN regions plays a crucial role in support of goal-directed WM processing while suppressing task-irrelevant internal thoughts and mind-wandering^28^. Evidence from dynamic functional interactions suggests that the SN is responsible for regulating a balance between FPN engagement and DMN disengagement to facilitate access to externally oriented stimuli and inhibit internally oriented attention during WM^29^. Unbalanced functional organization of these networks, with either hypo- or hyper-connectivity, has been seen in anxious individuals or under stress^12, 30^. However, how trait anxiety modulates the effects of long-term stress on functional balance of these networks during WM remains open.

Here we address the questions proposed above by leveraging fMRI and computational modeling of trial-by-trial decision responses to investigate how long-term stress and trait anxiety interact to affect dynamic decision computations during WM processing (**Fig. 1a**). In the long-term stress group, thirty-six
^*^ male participants were recruited from those who have been preparing for the upcoming competitive Chinese National Postgraduate Entrance Exam (CNPEE) for at least 6 months. Exposure to such an exam has been proven as a natural long-term psychosocial stressor by our and other laboratories^31, 32^. In the control group, thirty-two^†^ male participants matched in age and education who were not preparing for the CNPEE and did not have exposure to other major stressors in past 6 months were recruited. Participants underwent fMRI scanning while performing a numerical N-back task with a block design consisting of 0- and 2-back conditions (**Fig. 1b**). Trait anxiety and psychological distress were assessed by the State-Trait Anxiety Inventory^33^ and Symptom Checklist 90 (SCL-90)^34^ one day prior to the fMRI experiment. A Bayesian hierarchical version of the DDM (HDDM) was implemented to identify latent dynamic decisions during WM processing. Brain activation and network approaches were employed to identify how long-term stress and trait anxiety alter functional brain network balance during WM, including task-invoked activation/deactivation, as well as inter-network coupling amongst core nodes of the FPN, SN and DMN. Based on neurocognitive models of stress and anxiety, we expected that individual differences in trait anxiety would modulate the effects of long-term stress on latent dynamic decisions during WM, likely involving altered brain functional balance among the FPN, DMN, and SN regions at WM-related activation, deactivation, and network coupling levels.

**Fig 1.**
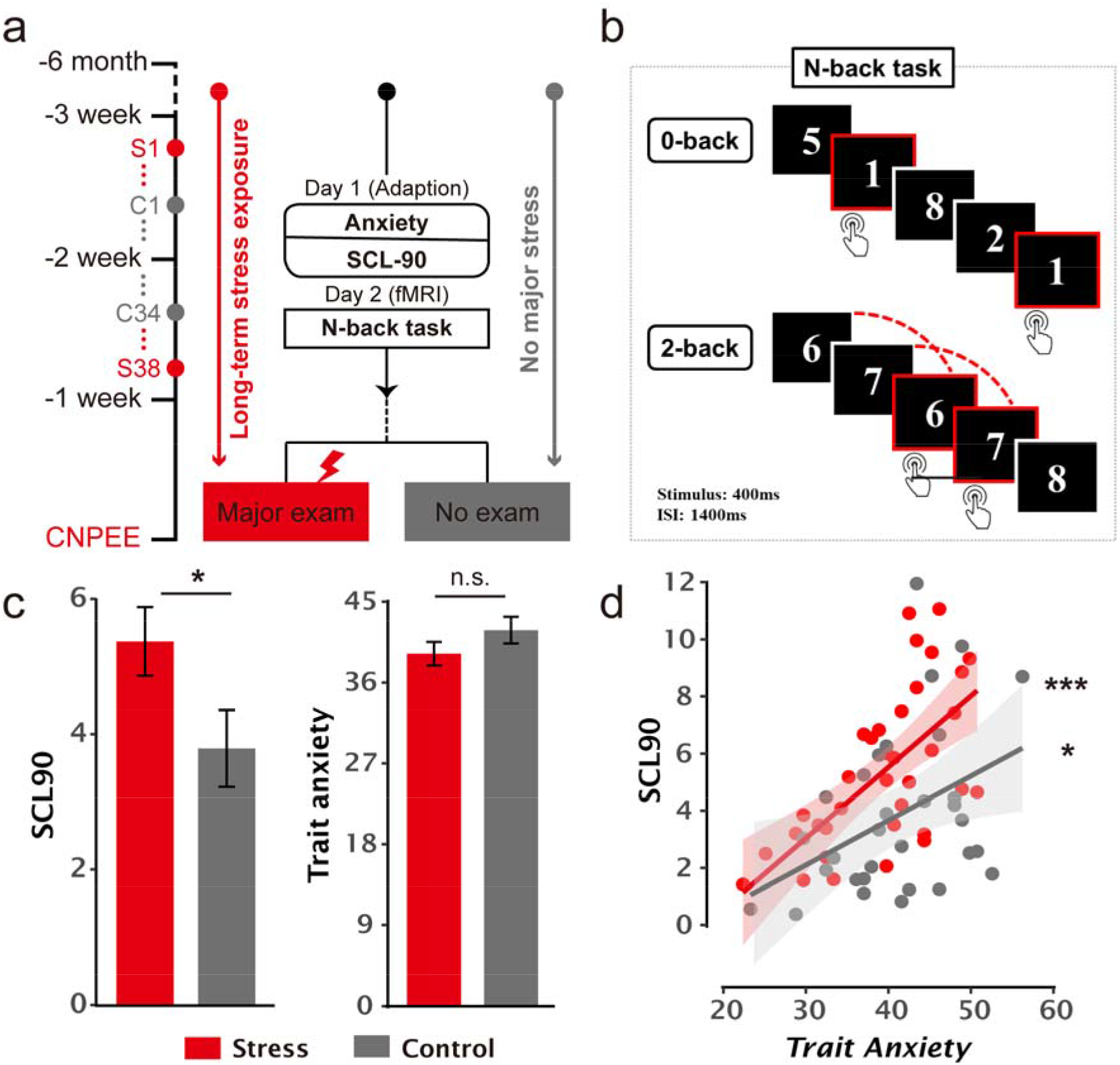
Experimental design and the effects of trait anxiety on psychological distress measurements amongst long-term stress and control groups. (**a**) An overview of experimental design illustrates the major procedures for participants from long-term stress and control groups with trait anxiety and SCL-90 assessments on the adaptation day (Day 1) prior to the fMRI N-back task (Day 2). These assessments occurred 1-3 weeks before the major exam stressor. There were 38 participants (S38) in the long-term tress group and 34 (C34) in the control group. Two participants from each group were excluded from further analyses due to excessive head motion during fMRI scanning, resulting in 36 and 32 participants in the stress and control groups, respectively (**b**) An illustration of the numerical N-back task that consists of 0- and 2-back conditions, with each digit item presented for 400 ms followed by an inter-stimulus interval of 1400 ms. Participants were instructed to detect whether the current item was ‘1’ in the 0-back condition, and were asked to decide whether the current item had appeared two positions back in the sequence in the 2-back condition. (**c**) Bar graphs depict psychological distress measured by SCL-90 and trait anxiety in the long-term stress and control groups. (**d**) Positive correlations of psychological distress with trait anxiety in long-term stress and control groups. (**b**) Psychological distress measured by Symptom Checklist 90 (SCL90) significantly differed between stress and control groups. (**c**) Positive correlation of psychological distress with trait anxiety in stress and control groups. (**d**) Trait anxiety scores in long-term stress and control groups. Notes: SCL-90, Symptom Checklist 90; \*\*\**P* < 0.001, \**P* < 0.05; n.s., not significant.

**Fig 2.**
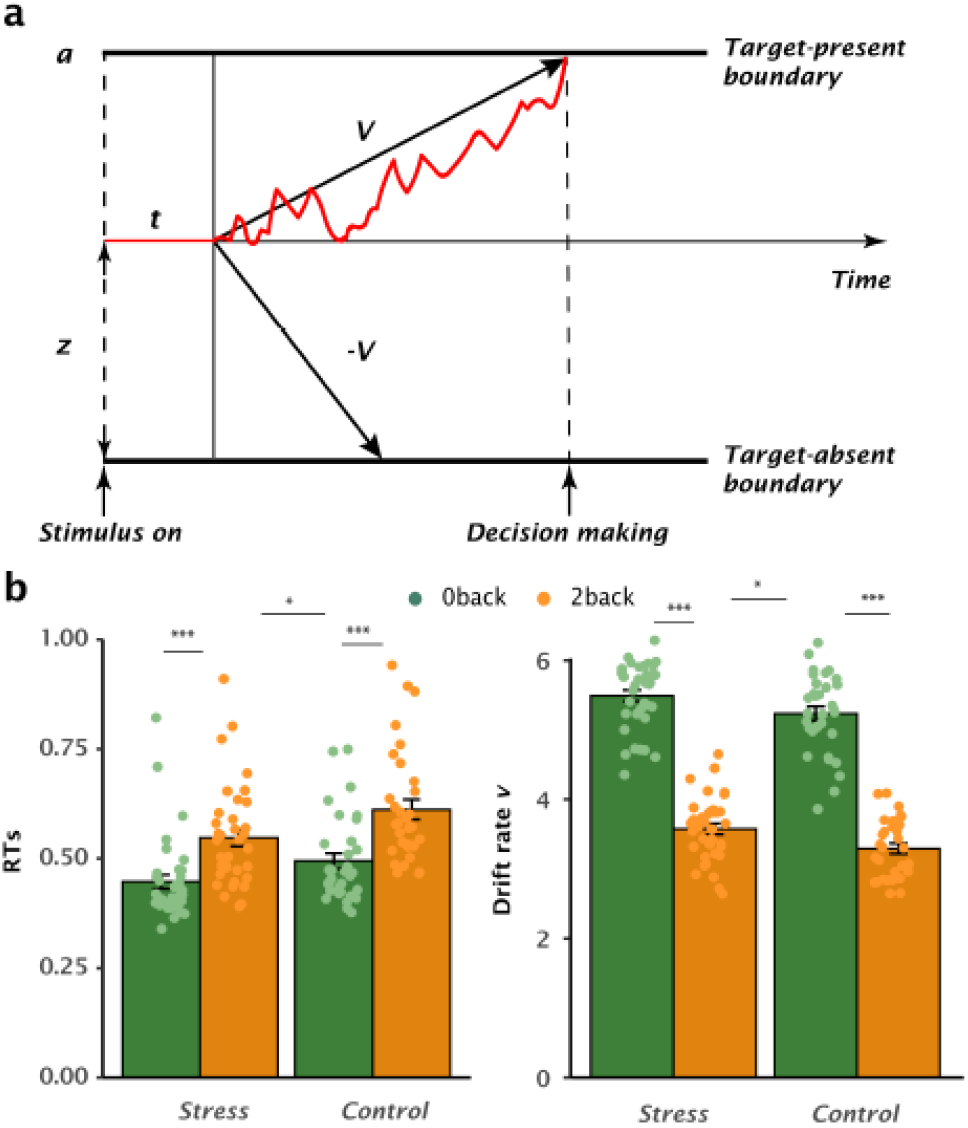
Effects of long-term stress on latent dynamic decision-making during WM. (**a**) Schematic view of the drift diffusion model (DDM) accounting for the n-back WM task with four model parameters: drift rate (*v*) indicates the rate of evidence accumulation until boundary threshold *a* is reached. The non-decision time *t* represents the time for stimulus encoding in addition to decision-making process. The starting point *z* reflects the prior preference toward one choice over the other. (**b**) Left panel: Mean reaction times (RTs) for 0- and 2- back conditions in the stress and control groups. Right panel: Estimated model parameter for drift rate in the DDM. Notes: Error bars represents SEM. Dots represent individual parameters.

## Results

### Effects of long-term stress and trait anxiety on psychological distress

We first investigated how long-term stress and trait anxiety affect psychological distress. Participants’ trait anxiety and psychological distress scores are listed in **Table S1**. No scores reached clinical levels of psychopathology based on SCL-90 norms^41^. Two-sample t-test revealed higher psychological distress in the stress than control group [*t(65) = 2.08, p = 0.041*](**Fig. 1a**). Moreover, individuals with higher trait anxiety exhibited greater psychological distress within both the stress [*r(34) = 0.65, p < 0.001*] and control groups [*r(29) = 0.41, p = 0.02*](**Fig. 1b**), even after regressing out state anxiety [stress: *r(34) = 0.48, p = 0.004*, control: *r(29) = 0.47, p = 0.008*]. Further analysis with Fisher’s z-transformed correlation coefficients revealed no group difference [z = 0.05, p = 0.96]. Prediction analyses based on machine learning using four-fold balanced cross-validation with linear regression confirmed that higher trait anxiety was predictive of greater distress after controlling for state anxiety in both groups (stress: r_(predicted, observed)_ = 0.46, *p* = 0.002; control: r_(predicted, observed)_ = 0.29, *p* = 0.036). These results indicate that individuals under long-term stress experience greater psychological distress than controls, with higher trait anxiety predictive of more distress in general.

### Effects of long-term stress on WM performance and latent dynamic decision measures

Next, we investigated how long-term stress affects WM performance and latent model-based measures. Separate 2-by-2 analyses of variance (ANOVAs) were conducted for accuracy and average RTs with Group (stress vs. control) as between-subject factor and WM load (0- vs. 2-back) as within-subject factor. These analyses revealed a main effect of WM load for accuracy and RTs [*both F(1, 66)* ≥ *5.21, p* ≤ *0.026*], with lower accuracy and slower RTs as task demands increased from 0- to 2-back condition [*t(66) =9.33, p < 0.0001*]. Critically, we also observed a main effect of Group for RTs [*F(1,66) = 5.37, p = 0.024*], which was mainly driven by faster RTs in the 2-back condition in the long-term stress group than controls [t(95.2) = -2.40, p = 0.018], though this was less pronounced in the 0-back condition [t(95.2) = -1.77, p = 0.08]. There was no main effect of Group on accuracy, nor Group-by-WM interactions for accuracy and RTs [all *F(1, 66) 2.57, p 0.11*].

We then investigated the effects of long-term stress on model-based measures during WM by fitting the HDDM to trial-by-trial RTs separately for 0- and 2-back conditions across participants. The model comparisons were performed for a total of 15 plausible model variants (**Fig. S4**). This yielded a model allowing for changes in parameters including drift rate *v*, decision threshold *a*, non-decision time *t*, and starting point *z* between conditions to provide the best fit and good convergence (**Table S3, Fig. S5**). Separate 2 (Group)-by-2 (WM-load) ANOVAs for model-based latent measures revealed a main effect of Group for drift rate [*F(1, 66) = 8.91, p = 0.004*] and decision-threshold [*F(1, 66) = 4.15, p = 0.046*]. Compared to controls, individuals under long-term stress exhibited faster drift rate [t(129) = 2.38, p = 0.019] with comparable decision threshold [t(83) = -1.44, p = 0.15] in the 2-back condition, and faster drift rate [t(129) = 2.17, *p* = 0.032] but less stringent threshold [t(83) = -2.39, *p* = 0.019] in the 0-back condition. The statistics for other measures are provided in **Table S4**. Both drift rate and decision threshold exhibited high correlations and the predictive ability to RTs in the 2-back condition (**Table S10-11**), indicating that model parameters can be recovered from the actual RTs. Together, these results indicate prominent effects of long-term stress on RTs during WM, along with faster drift rate mainly in moderate task demand but lower decision threshold in low task demand.

### Long-term stress shifts the balance between WM-related brain activation and deactivation

We further investigated how long-term stress affects brain systems during WM processing using whole-brain 2 (Group)-by-2 (WM-load) ANOVA. By contrasting the 2- with 0-back condition, we replicated robust WM-related activation and deactivation in core regions of the FPN and DMN respectively (whole-brain family wise error corrected *P* < 0.05) (**Fig 3a**). Importantly, a contrast reflecting the main effect of Group revealed a robust hyper-activation in the anterior insula (**Fig 3b**) and the middle occipital cortex extending into the cuneus (**Fig. S9**) in individuals under long-term stress, compared to than controls (**Table S12**) (voxel-wise *P* < 0.001, cluster *P* < 0.05 corrected). We also observed a Group-by-WM interaction in a set of distributed regions in the SN and DMN (**Fig. 4a-b**, **Table S13**). To verify whether these regions were engaged in WM-related (de)activation, we performed a conjunction analysis between the two contrasts reflecting Group-by-WM interaction and WM-related (de)activation (***Methods***). This revealed significant clusters in the dACC and the anterior insula (**Fig 4b**, **Table S13**) with greater WM-related activation in these regions in the long-term stress group than controls (voxel-wise *P* < 0.005, cluster *P* < 0.05 corrected). In the opposite contrast, we observed clusters in the DMN, with less WM-related deactivation in the medial prefrontal cortex (MPFC) and posterior cingulate cortex (PCC) in the long-term stress group than controls (**Fig 4a**, **Table S13**) (voxel-wise *P* < 0.005, cluster *P* < 0.05 corrected). These results indicate that long-term stress leads to hyper-activation in the SN regions, but less WM-deactivation in the DMN regions.

**Fig 3.**
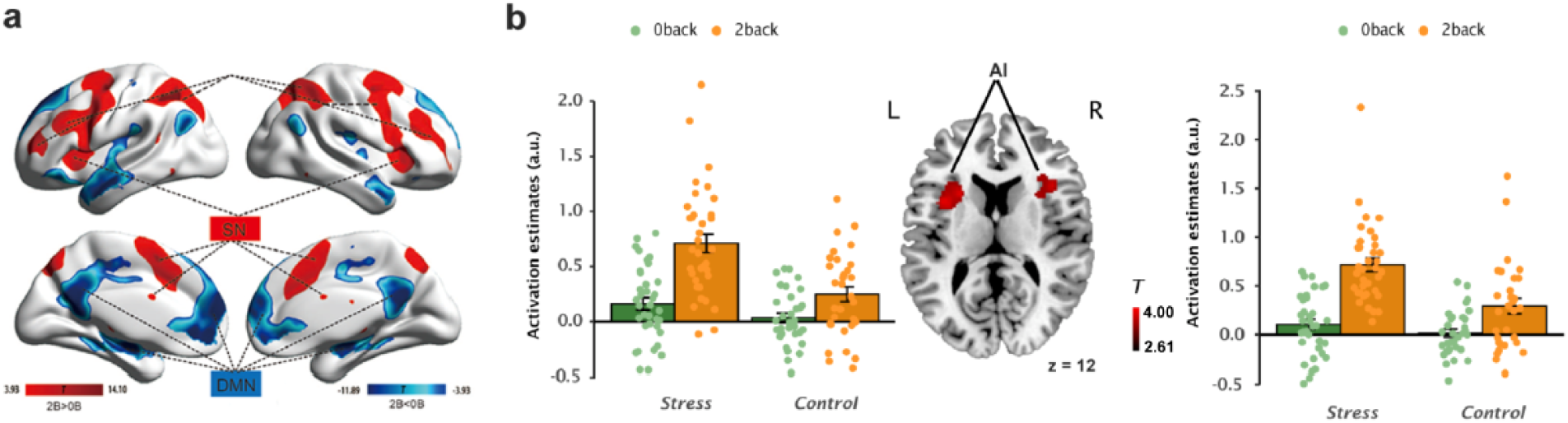
Brain regions showing main effects of WM load and long-term stress. (**a**) Significant clusters in distributed brain regions showing WM-related activation (red) and deactivation (blue) by contrasting 2- with 0-back conditions (whole-brain FWE *P* < 0.05 corrected). (**b**) Significant clusters in the bilateral anterior insula (middle) showing the main effect of long-term stress (voxel *P* < 0.001, and cluster *P* < 0.05 corrected) and corresponding parameter estimates extracted from the clusters. Color bar indicates minimum and maximal *T* values. Error bars represents standard error of the mean. Dots represent individual parameters. FWE, family wise error rate.

**Fig 4.**
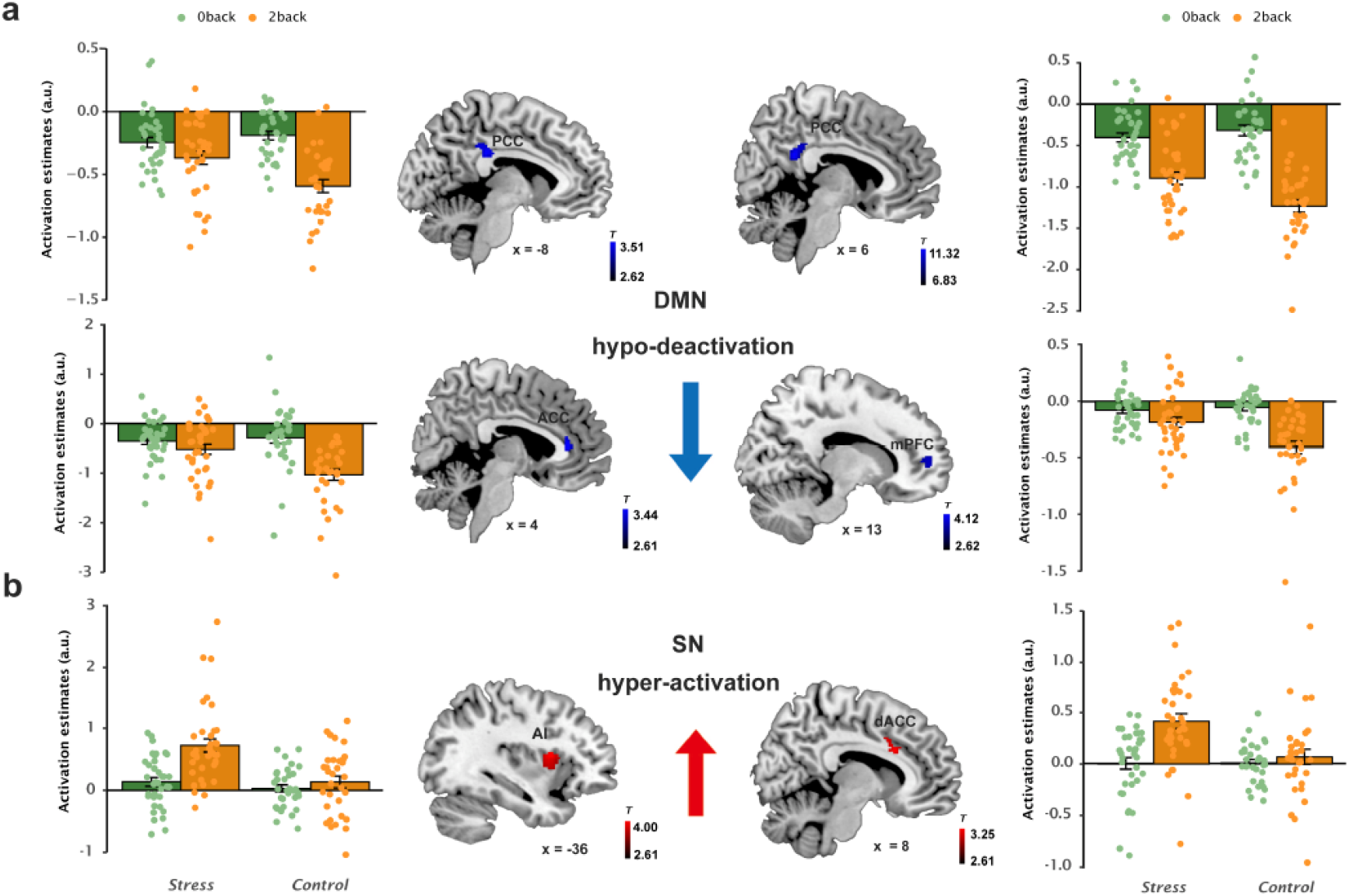
Long-term stress shifts the balance between WM-related brain activation and deactivation. The interaction effects between long-term stress and WM loads in brain regions of the default mode network (DMN) and salience network (SN). (**a**) Significant clusters in regions of the DMN including posterior cingulate cortex (PCC) and medial prefrontal cortex (mPFC) (middle panels), with weaker WM-related deactivation under long-term stress than control (voxel *P* < 0.005, and cluster *P* < 0.05 corrected). Bar graphs depict corresponding parameter estimates only for visualization purpose. (**b**) Significant clusters in regions of the SN including the dorsal anterior cingulate cortex (dACC) and anterior insula (AI) (middle panels), with greater WM-related activation under long-term stress than controls (voxel *P* < 0.005, cluster *P* < 0.05 corrected). Bar graphs depict corresponding parameter estimates only for visualization purposes. Color bars indicate *T* values which vary from image to image. Error bars represent the standard error of mean. Dots represent individual data points.

### Long-term stress and trait anxiety alter latent dynamic decisions through fronto-parietal activity

We then investigated whether trait anxiety modulates the effects of long-term stress on RTs and drift rate observed above. No direct associations of trait anxiety with RTs or drift rate were observed in either group (**Table S5**). Given the prominent effects of long-term stress on drift rate in the 2-back condition, we investigated how trait anxiety modulates neural correlates of drift rate in this condition under long-term stress compared to controls. Whole-brain independent sample t-test was conducted for WM-related activity maps between the two groups with drift rate as a covariate of interest. This analysis revealed a cluster in the IPS (voxel-wise *P* < 0.005, cluster *P* < 0.05 corrected), with higher WM-related activity in this region associated with slower drift rate in the long-term stress group [*r(34) = -0.48, p = 0.003*], but an opposite pattern in the control group [*r(30) = 0.40, p = 0.024*](**Fig 5a**). Further analysis for Fisher’s z-transformed correlations revealed a group difference [z = -3.71, *p* < 0.001]. When restricting our analysis to the stress group (see Methods for the rationale), we also observed a significant cluster in the MFG (voxel-wise *P* < 0.005, cluster *P* < 0.05 corrected), with higher WM-related activity in this region associated with slower drift rate under long-term stress (r(34) = -0.52, p = 0.001), but no reliable correlation in controls (r(30) = 0.19, p = 0.31) (group difference: z = -3.02, p = 0.003). Prediction analyses confirmed that higher activity in the IPS and MFG was predictive of slower drift rate in the long-term stress group (**Table S11**). Parallel analyses for decision threshold revealed no reliable correlation with WM-related brain activity.

**Fig 5.**
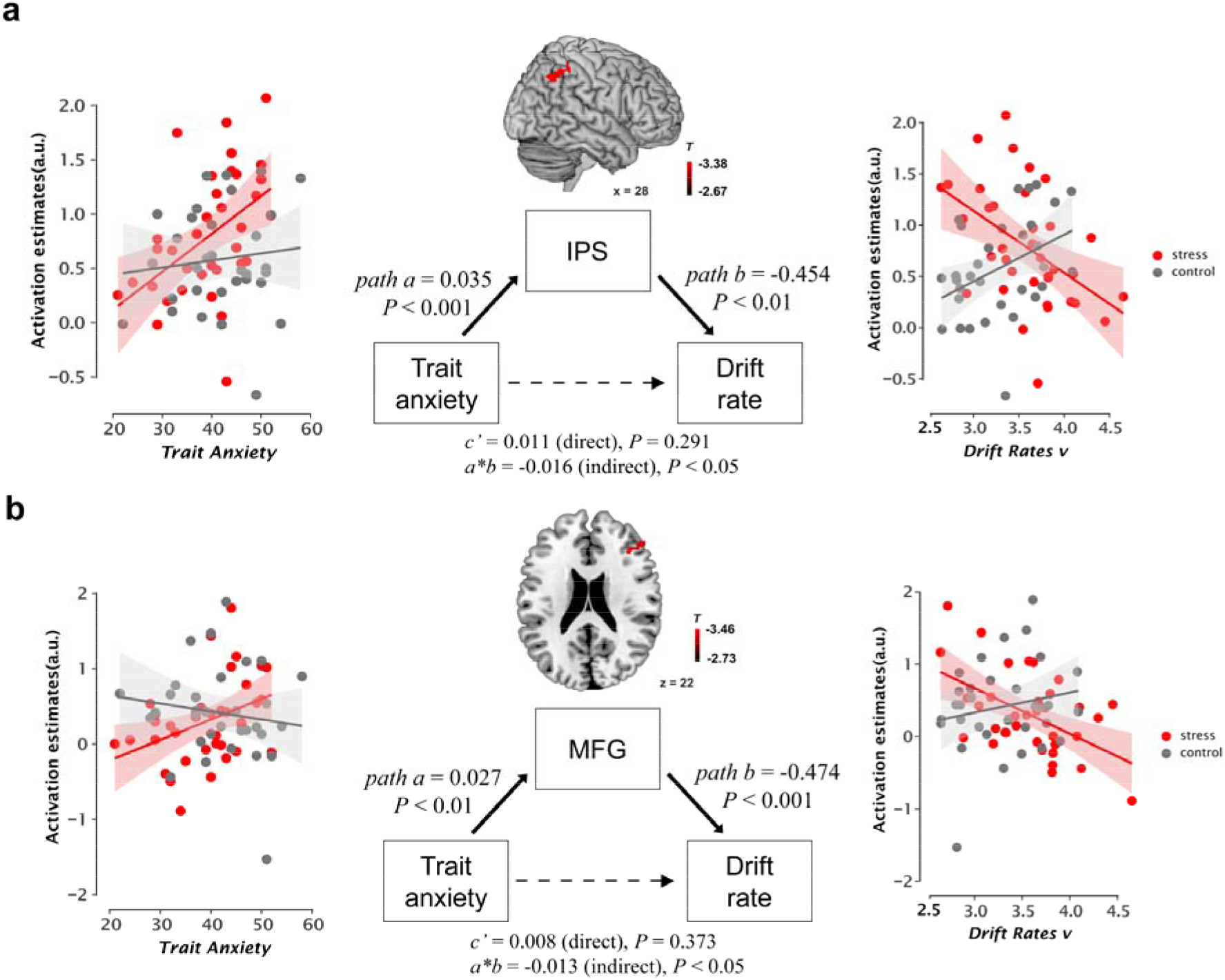
The relation between trait anxiety, brain activity, and drift rate. (**a**) Left and right panels: Scatter plots depict the correlations of individual’s trait anxiety with WM-related parietal activity that in turn were correlated with drift rate in the long-term stress and control groups. Middle panel: The mediating effect of WM-related activity in the intra parietal sulcus (IPS) on the relationship between trait anxiety and drift rate (voxel *P* < 0.005, cluster *P* < 0.05). (**b**) Left and right panels: Scatter plots depict the correlations of individual’s trait anxiety with WM-related prefrontal activity that in turn were correlated with drift rate in the long-term stress and control groups. Middle panel: The mediating effect of WM-related activity in the middle frontal gyrus (MFG) on the relationship between trait anxiety and drift rate (voxel *P* < 0.005, cluster *P* < 0.05). Color bar indicates minimum and maximal T values. Notes: \**P* < 0.05, \*\**P* < 0.005, \*\*\**P* < 0.001.

We then investigated whether trait anxiety modulates the correlations between WM-related fronto-parietal activity and drift rate under long-term stress in comparison to controls. Although drift rate displayed no reliable correlation with trait anxiety in either group [*r ≥ -0.09, p ≥ 0.60*], we observed positive correlations of trait anxiety with WM-related fronto-parietal activity in the long-term stress group (IPS: r(34) = 0.48, p = 0.003; MFG: r(34) = 0.39, p = 0.02) but not controls (IPS: r(30) = 0.11, p = 0.54; MFG: r(30) = -0.14, p = 0.44) (**Fig 5a&b**). Prediction analyses confirmed that higher trait anxiety was predictive of higher activity in the IPS and MFG (**Table S11**). Further tests revealed a group difference in correlations for the MFG (z = 2.16, p = 0.03) but not IPS (z = 1.62, p = 0.11).

Given higher activity in the IPS and MFG were reliably associated with higher trait anxiety and slower drift rate in the long-term stress group, one may conjuncture that WM-related activity in these regions could act as a mediator to account for an indirect association between trait anxiety and drift rate. We thus implemented multi-group structural equation models (SEMs) to test potential mediation effects of WM-related activity in the IPS and MFG on the indirect associations between trait anxiety and drift rate in the long-term stress and control groups. These analyses revealed significant mediation effects of WM-related frontoparietal activity *[IPS: indirect Est. = -0.016, 95%CI = [-0.034, -0.006]; MFG: indirect Est. = -0.013, 95%CI = [-0.029, -0.003]* on the indirect association between trait anxiety and drift rate under long-term stress (**Fig 5a&b**) but not controls (**Fig S7**, **Table S7**). Critically, further analyses revealed significant group differences in the mediation effects for the IPS (*Est. = -0.018, 95%CI = [-0.038, -0.005]) and MFG (Est. = -0.012, 95%CI = [-0.028, -0.001]*). Parallel analyses for state anxiety, however, exhibited no group differences in the mediation effects for the IPS (*Est. = -0.007, 95%CI = [-0.020, 0.006]*) and MFG (*Est. = -0.008, 95%CI = [-0.025, 0.003]*) between long-term stress and controls (**Table S8**). Together, these results indicate that individuals with higher trait anxiety are prone to exhibit slower evidence accumulation than controls mediated through higher frontal-parietal activity during WM under long-term stress.

### Long-term stress and trait anxiety alter large-scale functional brain network balance during WM

To further investigate how long-term stress and trait anxiety affect functional coordination of large-scale brain networks during WM decision processing, we analyzed intra- and inter-network coupling and decoupling among the FPN, DMN and SN regions in the long-term stress group and controls. The FPN, DMN and SN nodes were independently defined to avoid double dipping or selection biases, then intra- and inter-network coupling metrics were computed for each participant (**Fig 6a**). Separate 2 (Group)-by-2 (WM-load) ANOVAs were conducted for these network data. For intra-network coupling, we found a main effect of WM-load for the FPN [*F(1, 66) = 4.17, p = 0.045*], but no long-term stress effects nor Group-by-WM interactions were found [*all F(1,66)* ≥ *0.78, p > 0.10*] (**Table S21**).

**Fig 6.**
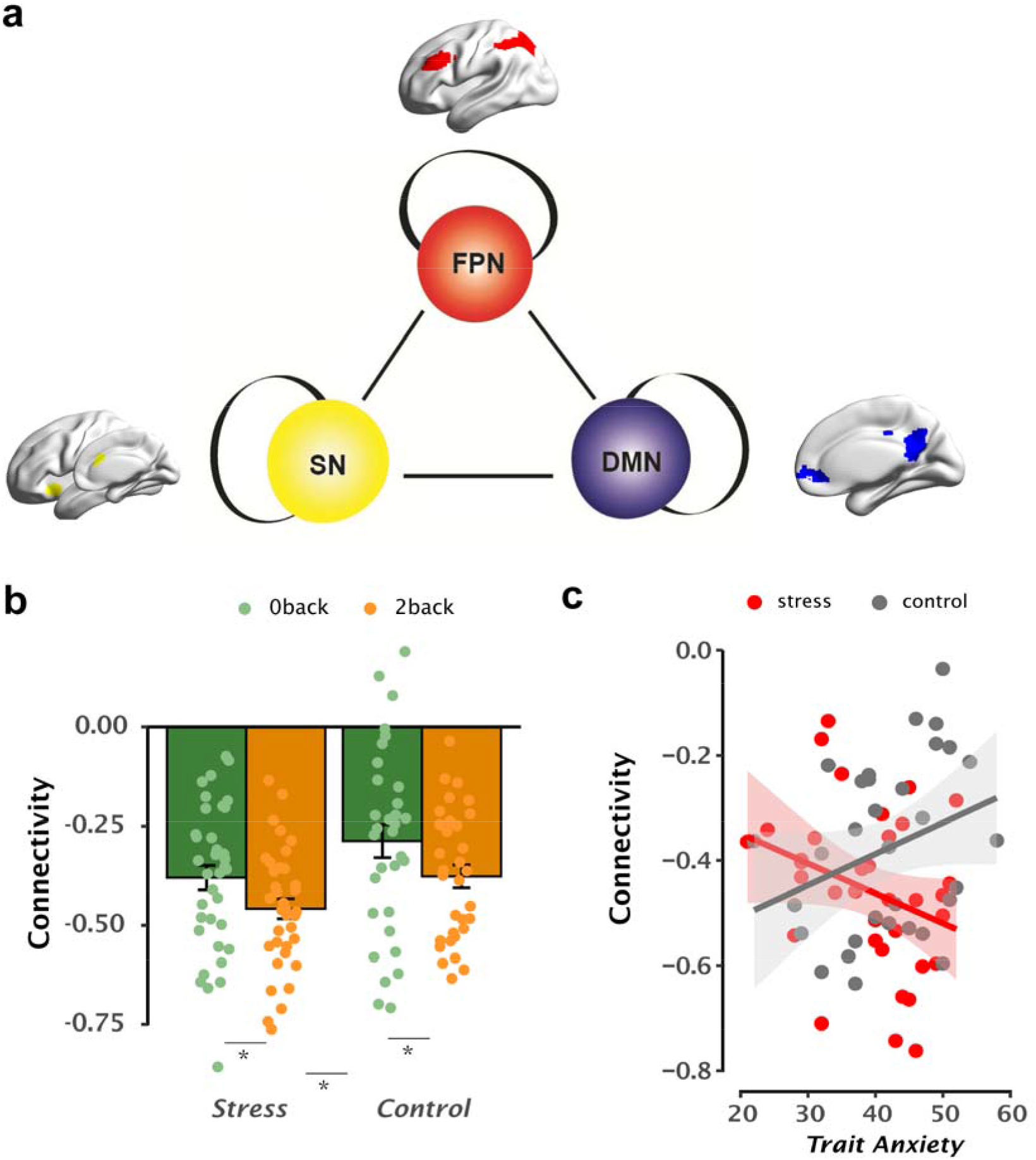
Inter-network connectivity modulated by trait anxiety under long-term stress. (**a**) Representative nodes of the three core brain networks involved in WM processing, including the fronto-parietal network (FPN), default mode network (DMN) and salience network (SN). (**b**) Bar graphs depict the main effect of long-term stress on inter-network coupling between the FPN and DMN in 0-and 2-back conditions, with greater FPN-DMN decoupling in the long-term stress group in comparison to controls. (**c**) Scatter plot depicts an interaction effect between long-term stress and trait anxiety on FPN-DMN decoupling, with a negative correlation between decoupling strength and trait anxiety in the long-term stress group, but an opposite pattern in the control group. Solid lines represent the average, and shaded areas represent 95% confidence intervals. Dots represent individual data points.

For inter-network coupling, we found a main effect of WM-load for both FPN-DMN decoupling [*F(1,66) = 12.79, p < 0.001*] and SN-DMN coupling [*F(1,66) = 4.67, p = 0.034*] with increased decoupling in the 2- than 0-back condition (all *t(66) > 3.00, p < 0.01*). Moreover, we found a main effect of Group [*F(1, 66) = 3.36, p = 0.028*] only for FPN-DMN decoupling, with greater decoupling under long-term stress than controls [*t(66) = -2.25, p = 0.028* (**Fig 6b**). However, no Group-by-WM interaction was observed (**Table S21**). Control analyses using nodes from meta-analysis of previous studies yielded similar effects (**Fig. S8**). Critically, individuals with higher trait anxiety exhibited stronger FPN-DMN decoupling under long-term stress [*r(34) = -0.36, p = 0.034*], but an opposite pattern was found in controls *[r = 0.32, p = 0.079]* in the 2-back condition even after controlling for state anxiety (**Fig 6c**). Prediction analyses confirmed that higher trait anxiety was predictive of stronger FPN-DMN decoupling under long-term stress (**Table S11**). Further analysis revealed a significant group difference [z = -2.78, p = 0.005], indicating a prominent interaction between trait anxiety and long-term stress on FPN-DMN decoupling. Interestingly, higher FPN-DMN decoupling was associated with lower drift rate in the stress group [*r(34) = 2.19, p = 0.036*]. No mediation effects, however, were observed among trait anxiety, FPN-DMN decoupling, and drift rate in either group. These results indicate that long-term stress leads to increased FPN-DMN decoupling, and higher trait anxiety is predictive of stronger FPN-DMN decoupling in those under long-term stress but not in controls.

## Discussion

In this study, we investigated the neurocognitive mechanisms of how long-term stress and trait anxiety interact to affect dynamic decision computations during WM. We found that long-term stress led to higher psychological distress, faster RTs, and faster drift rate, but a lower decision-threshold than controls, with higher trait anxiety predictive of greater distress. These effects occurred with general hyper-activation in the anterior insula, greater WM-related activation in SN regions, and less WM-related deactivation in DMN regions. Moreover, individuals with higher trait anxiety were prone to slower drift rate via higher WM-related activity in FPN regions in the long-term stress but not control group. Long-term stress also led to stronger DMN decoupling with the FPN than controls, with higher trait anxiety predictive of stronger FPN-DMN decoupling in those under long-term stress. Our findings provide a neurocognitive account for the interplay of long-term stress and trait anxiety on latent dynamic decisions during WM, via altered functional brain network balance among FPN, DMN and SN regions.

As expected, individuals in the long-term stress group experienced higher psychological distress than controls, indicating the effectiveness of our natural long-term stress paradigm. Moreover, individuals with higher trait anxiety experienced more psychological distress in general, even after controlling for state anxiety. These results are in line with previous findings on sustained distress and other symptoms in chronic stress^35^, which agrees with the psychological view of trait anxiety as a vulnerable phenotype of stress-related psychopathology^7, 8^. Behaviorally, individuals under long-term stress exhibited faster RTs but comparable accuracy during WM than those in controls. Higher drift rate in the 2-back condition but a less stringent decision threshold in the 0-back condition was further observed by computational modeling of trial-by-trial decisive responses. In accordance with integrative models of stress, anxiety, and cognitive performance^9, 36^, our results show that sustained exposure to exam stress may not impair performance effectiveness (i.e., comparable accuracy), and may enhance processing efficiency (i.e., faster RTs and drift rate) under moderate task demand conditions. However, enhanced efficiency differs from previously reported cognitive deficits of chronic stress^2, 3^. Two factors are critical to reconcile this discrepancy. First, according to the Yerkes–Dodson law, the effects of stress on behavioral performance exhibits a nonlinear inverted-U shape curve as a function of stress severity and task difficulty. Thus a beneficial effect can be reached at moderate levels of stress and task demands^37^. In this view, our observed faster RTs and drift rate may reflect enhanced processing efficiency at moderate WM-load in those under exam stress. Likewise, one previous study reported that stressed participants reacted faster at moderate WM-load^38^. Additionally, our observed less stringent decision threshold under low task demand may reflect stress-induced tendency to a liberal response bias^39^.

Another factor is the interplay of long-term stress with trait anxiety that entails an individual’s resilience and vulnerability to maladaptation^7, 40^. When taking individual’s trait anxiety into account, lower trait-anxious individuals exhibited relatively faster drift rate in those under long-term stress than in controls (Figure S6), while accuracy remained at a comparable level across groups and anxiety levels. In other words, stress-induced faster drift rate is driven by low-trait anxious individuals, suggesting that a beneficial form of adaptation to sustained exam stress is likely driven by enhanced processing efficiency in those individuals. This is consistent with previous studies on stress vulnerability reporting that some individuals seem to act as “resilient” agents who can develop adaptive strategies to cope with stress^6, 7, 40^. Furthermore, our observations on high trait-anxious individuals may reflect the recruitment of compensatory strategies to prevent shortfalls in accuracy according to an influential model of cognitive trait anxiety and performance^41^. As we discuss later, the interplay of long-term stress and trait anxiety on drift rate was further modulated by a shift in brain functional balance at WM-related (de)activation and network (de)coupling levels.

At the brain activation level, individuals under long-term stress exhibited a general hyper-activation in the anterior insula regardless of WM load. Similar hyper-activation was reported by previous studies on hyper-vigilance in those experiencing distress, as well as in anxious individuals in healthy and psychiatric populations^42^. Thus, our observed hyper-activation may reflect an increase in emotional awareness of distressed feelings^43^. Similar hyper-activation has been linked to increased sensitivity to sensorimotor processing, which could explain the aforementioned liberal response bias under long-term stress. Moreover, long-term stress led to greater WM-related activation in core nodes of the SN including the anterior insula and dACC, but less WM-related deactivation in the DMN regions. The anterior insula and dACC are thought to support salience processing, emotional awareness of distress, and executive function^25, 44^. Based on the dual competition model of emotion-cognition interaction^45^, such SN regions are responsible for reallocating neural resources to resolve competition between emotional processing and executive function. Thus, greater SN engagement under long-term stress may reflect recruitment of additional effort to cope with stress reactivity and to regulate stress-induced distress feelings along with related thoughts that are irrelevant to the WM task. Stress-induced task-irrelevant internal thoughts were indicated by accompanying less DMN deactivation, which parallels the empirical findings of aberrant DMN suppression in psychiatric diseases such as depression^46^. It is possible that our observed less WM-related DMN deactivation may interact with greater SN activation to accomplish the WM task while suppressing distress feelings and irrelevant thoughts.

With respect to long-term stress and trait anxiety interactions on brain-behavior relationships, individuals with higher trait anxiety under long-term stress exhibited an indirect association with slower drift rate through higher WM-related frontoparietal activity but not those in controls. The lack of a mediation effect in the control group suggests that the FPN serves as a mediator only in individuals under long-term stress. Such mediation effects parallel the cognitive models of anxiety and related studies showing that high trait anxiety impairs processing efficiency but not performance effectiveness on tasks involving executive function under stressful conditions^9^. As discussed earlier, anxious individuals may recruit additional resources as a compensatory strategy to achieve comparable performance effectiveness^10, 11, 41^. Given the dlPFC and IPS have been associated with drift rate in the process of evidence accumulation in human and non-human primate studies with single cell recording^17, 18, 47^, our observed higher frontoparietal engagement provides neuroimaging evidence to suggest that higher trait-anxious individuals under long-term stress tend to recruit more neural resources to maintain comparable performance at the cost of the speed of evidence accumulation to make correct decisions. Specifically, when performing the 2-back task under a normative condition, one must constantly update and maintain the most recent 2-items in mind and accumulate sufficient evidence extracted from each rapidly presented stimulus to ensure a correct decision whether the current item is a target or not^48^. Under stress, however, high trait-anxious individuals experienced more distress likely with a hypervigilant state as indicated by hyper-activation in the SN. This might have impeded efficient extraction of target-relevant information due to potential confounds by feelings of distress, irrelevant thoughts, or other noise. Hence, more time is needed to accumulate sufficient evidence to reach a decision, thereby slowing the speed of evidence accumulation. Indeed, recent studies reported that the negative impact of trait anxiety extends beyond aversive feelings and involves impediment of ongoing goal-directed behaviors. This then results in an impaired capacity to disengage from the previously relevant sensory information to overcome distracting stimuli^12, 41^.

According to the neurobiological models of stress, the major targets of stress-sensitive hormones include regions of the FPN critical for drift rate during WM processing^49^. Given the link of high trait anxiety to stress sensitivity and stress hormone release^8^, we thus speculate that excessive stress hormones might in part account for the higher activity in the IPS and MFG in trait-anxious individuals under long term-stress. By extending such neurobiological accounts, our findings provide new insights suggesting that high trait anxiety per se does not necessarily lead to cognitive deficits. Rather, high trait anxiety works in concert with long-term stress to determine the (mal)adaptive effects on human brain and cognition. Together, under long-term stress, a slower speed of evidence accumulation in higher trait-anxious individuals may reflect less efficient evidence accumulation in the process of dynamic decisions during WM, likely via increased FPN engagement in order to make correct responses.

At the brain network level, we found greater FPN-DMN decoupling during WM under long-term stress than controls. Empirical evidence from previous studies has demonstrated that such increased decoupling likely reflects more effort to suppress task-irrelevant internal thoughts and mind wandering^50^. Likewise, stronger FPN-DMN decoupling here may reflect recruitment of additional effort to suppress task-irrelevant internal thoughts, while performing the goal-directed WM task. This notion is also in line with our observed faster RTs and drift rate under long-term stress. Critically, individuals with higher trait anxiety exhibited stronger decoupling between DMN and FPN regions in those under long-term stress but not in controls, and such stronger decoupling was then associated with slower drift rate. This again provides evidence to suggest that high trait-anxious individuals might recruit additional resources relying on FPN-DMN decoupling, along with the elevated FPN activity mentioned above, to make correct responses and prevent a shortfall in accuracy at the cost of processing efficiency. Notably, the SN, especially the anterior insula and dACC, is thought to play a role in generating control signals to regulate switching between the FPN engagement in goal-directed tasks and DMN disengagement from irrelevant thoughts and mind wandering^29, 44, 51^. Although we did not find effects of long-term stress and/or trait anxiety on SN coupling with other networks, this switching mechanism is still relevant to account for our observed hyper-activation of the SN, along with less WM-related deactivation in the DMN and the increased DMN-FPN decoupling under long-term stress. Such alterations in functional brain network balance may reflect shifted attention out of internally-driven mental activity (i.e., stress-related distress feelings) in order to make correct decision during a WM task.

It is worth noting that although trait and state anxiety are recognized as two distinct constructs in psychometric theory, they are inherently inter-correlated and thus challenging to dissociate. Nevertheless, our conclusions on trait anxiety still hold after controlling for state anxiety. Notably, the mediation effects of WM-related fronto-parietal activity on an indirect association between high trait anxiety and slower drift rate are only present in those under long-term stress. Such mediation effects are not present for those with higher state anxiety. The following limitations should be considered. First, we only included male participants to mitigate potential confounds related to menstrual cycles^52^, which limits the generalizability of our findings. Second, individual’s intelligence may complicate our observed effects of long-term stress and trait anxiety on WM, though null effects of long-term stress or trait anxiety on WM accuracy may neutralize this concern. Third, individuals exposed to long-term exam stress might experience sleep disruption and other stressors that could complicate our findings. Fourth, our block design for the WM task precludes trial-by-trial parametric modulation analyses linked to computational measures. Because there are a relatively small number of trials in our WM task, we used the HDDM given its suitability for estimating model parameters with few trials across participants^53^. In fact, our validation analyses showed a good model fit as reported by previous studies^54, 55^, indicating that model parameters can be reliably recovered from actual RTs. Future studies with novel designs are needed to resolve these limitations.

**In conclusion**, our study demonstrates that long-term stress and trait anxiety interplay to affect latent dynamic decisions during WM by altering brain network balance in core regions of the SN, FPN, and DMN. Our findings point toward a neurocognitive model suggesting that trait anxiety modulates latent decision-making dynamics during WM under long-term stress. This may inform personalized assessments and preventions for stress-related (mal)adaptation.

## Methods

### Participants

Seventy-two healthy male senior college students participated in this study. Thirty-eight participants (age range: 20-24 years old, mean ± S.D. = 21.57 ± 0.83) in the long-term stress group were recruited 1-3 weeks before a highly competitive Chinese National Postgraduate Entrance Examination (CNPEE). An independent cohort of 34 male participants matched in age and education (age range: 20-24 years old, mean ± S.D. = 21.61 ± 0.92) who did not participate in the CNPEE or have any other anticipated stressor were recruited to the control group. Inclusion criteria for long-term exam stress were as follows (Figure S1): (1) Participants had been preparing for the upcoming competitive CNPEE for at least 6 months. (2) Participants had to provide the CNPEE certificate registered more than 6 months before the experiment. (3) They had to participate the experiment within a 1 to 3-week time window before the CNPEE to ensure that they were experiencing high levels of psychosocial stress. We didn’t included female participants in order to mitigate potential confounds of their menstrual cycles^52^. Informed written consent was obtained from all participants prior to the experiment, and the study protocol was approved by the Institutional Review Board for Human Subjects at Beijing Normal University. Four participants (two in each group) were excluded from further analyses due to head movement more than one voxel in translation or in rotation.

### General experimental procedure and working memory (WM) task

Both psychological distress and trait anxiety measures were administrated one day before the fMRI experiment to mitigate potential confounding effects for their self-reports that may suffer from bias during times of acute stress in the task experiment. On the experimental day, participants were instructed to practice the task before fMRI scanning^56^. We used the 0- and 2-back conditions only, in order to create a robust contrast between low and moderate task demands to gain the differences in modeling parameters, brain activation, and connectivity measures between these two conditions. Participants then underwent fMRI scanning while performing the N-back task.

The entire N-back task included ten blocks, which alternated between five 0-back blocks and five 2-back blocks, interleaved by a jittered fixation that ranged from 8 to 12 sec. Each “block” refers to an experimental condition that consists of a sequence of items presented continuously for an extended time interval to maintain a steady state of certain cognitive processing such as 0- or 2-back condition here. In each block, a pseudorandomized sequence that included 15 digits was presented, with a 400 ms duration of each digit, followed by an inter-stimulus-interval of 1400 ms. Each block started with a cue indicating the 0-back or 2-back condition. In the 0-back task, participants were instructed to detect whether the current item was ‘1’. In the 2-back task, participants were asked to decide whether the current item had appeared two positions back in the sequence (**Figure 1b**). When detecting a target, participants were required to press a button with their index finger as quickly and accurately as possible, and to withhold their response for target-absent trials. There were 21 targets and 17 targets for the 0- and 2-back conditions respectively.

### Psychological measurements of long-term stress

The State-Trait Anxiety Inventory^33^, one of the most commonly used scales to measure trait anxiety level in healthy populations, was collected both in the stress and control groups. The Symptom Checklist (SCL-90) was used to evaluate psychological distress including symptoms of psychopathology under long term stress^57^. One outlier larger than 2.5 standard deviations from the control group was excluded for SCL data.

### Drift diffusion modeling for trial-by-trial decision responses in WM

The DDM conceptualizes decision-making as an evidence accumulation process in which effective evidence is extracted from the representations of stimuli that are inherently variable and noisy, and gradually accumulated over time until sufficient evidence reaches the decision threshold and a choice is executed (**Fig 2a**). For the 2-back task, such evidence accumulation process can be considered as accumulating effective information in mind on the exact position of each item in order to make a precise decision whether the current item appeared two positions back in the sequence. The DDM was then implemented to decompose participants’ trial-by-trial RTs during WM into latent processes which were modulated by the following free parameters: 1) drift rate *v* reflects the speed of evidence accumulation which depends on the presented stimulus and task difficulty; 2) decision threshold *a* determines how much evidence has to be accumulated until a decision in made and thus reflects the level of cautiousness; 3) non-decision time *t* reflects processes unrelated to the decision (e.g., sensory processing in visual areas and motor execution of the choice); 4) starting point *z* reflects the prior response bias or preference toward one choice over the other^58^.

The DDM is known as a de-facto standard for the two-alternative forced choice tasks, in which the two choices correspond to the upper and lower decision boundaries respectively^59^. Recent advances have extended the DDM to many task paradigms with one choice such as Go/no-Go task, as RTs of no-Go condition cannot be measured^60, 61^. Likewise, we fitted the DDM to trial-by-trial RTs for hits (target items with successful response) and false alarms (non-target trials with response) in 0- and 2-back conditions. The DDM parameters were then estimated by the HDDM across participants for its suitability to a relatively small number of trials, according to the most recent simulation data by systematic comparisons of multiple drift models^53^. Critically, the hierarchical modeling formulated in the Bayesian framework allows us to simultaneously estimate parameters on both group and individual levels, in a way that individual parameters were drawn from the group distribution^58^. Differences in RTs between 0- and 2-back conditions, and between stress and control groups, implicate changes in one or more DDM parameters between task conditions and groups. To examine whether the four parameters varying 0- and 2-back conditions led to greater biases between different models, 15 variants of the DDM with different parameter constraints were established for both stress and control groups (Figure S4). Model comparisons were conducted by using Deviance Information Criterion (DIC)^58^. For each model, Markov chain Monte Carlo (MCMC) sampling methods were applied to perform Bayesian inference by generating 20000 samples and discarding the first 2000 samples as burn-in. The best model is determined by the minimum DIC. We further employed the Gelman-Rubin statistic to assess the convergence of the model. Note that a difference of 10 in DIC is considered acceptable^62^. The value of computed for all parameters were close to 1.0 and less than 1.01, indicating good convergence where successful convergence is indicated by values < 1.1^58^. The four parameters of each participant from the best fitted model were then submitted to subsequent analyses.

### Behavioral data analysis

Two-sample t-tests were conducted to compare the differences in trait anxiety and psychological distress between groups. Correlation analyses were conducted to compute the relationships of trait anxiety with psychological distress and latent dynamic decision measures from HDDM. Statistical tests were conducted to compare group differences in Fisher r-to-z transformed correlation coefficients. Separate AVNOAs were conducted to examine the effects of long-term stress on conventional and latent computational measures for the 0-back and 2-back conditions. Mixed factorial ANOVAs were conducted using the “afex” R package^63^, and the Greenhouse-Geisser correction was applied whenever a non-sphericity assumption was violated.

### Imaging data acquisition

Participants were scanned in a Siemens 3.0-Tesla TRIO MRI scanner (Erlangen, Germany) at the Brain Imaging Center of the National Key Laboratory of Cognitive Neuroscience and Learning at Beijing Normal University. Functional images were acquired with a gradient-recalled echo planar imaging sequence (axial slices 33, repetition time 2000 ms, echo time 30 ms, flip angle 90°, slice thickness 4 mm, gap 0.6 mm, field of view 200 × 200 mm, and voxel size 3.1 × 3.1 × 4.6 mm^3^). Functional imaging session lasted 464 seconds during the N-back WM task. To improve individual coregistration and spatial normalization, a high-resolution anatomical image was acquired in the sagittal orientation using a T1-weighted 3D magnetization-prepared rapid gradient echo sequence (slices 192, repetition time 2530 ms, echo time 3.45 ms, flip angle 7°, slice thickness 1 mm, field of view 256 × 256 mm, and voxel size 1 × 1 × 1 mm^3^).

### Imaging data analysis

#### Preprocessing

Imaging data analysis was performed using Statistical Parametric Mapping 8 (SPM8 https://www.fil.ion.ucl.ac.uk/spm/software/spm8/). The first four functional volumes were discarded to enable T1 equilibration. The remaining volumes were first realigned to correct for head motion. The realigned volumes were then corrected for slice acquisition timing. The mean functional image was coregistered to each participant’s T1-weighted structural image and then normalized to a standard stereotaxic Montreal Neurological Institute (MNI) space with a resolution of 2 × 2 × 2 ^3^ . The mm functional images were then spatially smoothed by an isotropic Gaussian kernel with 6-mm full-width at half-maximum.

#### Statistical analysis

Smoothed images were statistically analyzed under the general linear model (GLM) framework in SPM8. To assess neural activity associated with 0-back and 2-back conditions, these conditions were modeled separately as boxcar regressors and convolved with the canonical hemodynamic response function built in SPM8. Additionally, six realignment parameters from preprocessing were included to account for movement-related variability. The analysis included high-pass filtering using a cutoff of 1/128 Hz and a serial correlation correction using a first-order autoregressive model (AR[1]).

Corresponding contrast parameter images for 0- and 2-back conditions at the individual level were then submitted to a second-level group analysis using 2-by-2 factorial ANOVA, with Group (long-term stress vs. control) as between-subject factor and WM-load (0- vs. 2-back) as within-subject factor to examine the main effects of Group and WM-load, and their interaction on task-invoked brain response. We identified brain regions showing significant Group and WM-by-Group interaction effects, and then applied a conjunction analysis of the minimum statistic^64^ with the contrasts of ‘2- > 0-back’ and ‘0- > 2-back’ separately. This allows us to identify brain regions commonly showing WM-by-Group interaction and WM-related activation/ deactivation. Significant clusters were determined by a voxel-wise height threshold of *p < 0.001* and an extent threshold of *p* < 0.05 corrected for multiple comparisons using suprathreshold cluster-size approach based on Monte-Carlo simulations. Given our priori hypotheses regarding the DMN, SN and FPN regions, these regions were additionally investigated using a height threshold of *p < 0.005* and an extent threshold of *p < 0.05* corrected for multiple comparisons. Monte-Carlo simulations were implemented using the AlphaSim procedure. Ten thousand iterations of random 3D images, with the same resolution, dimensions and 6-mm smoothing kernel as used our fMRI data analysis, were generated. The maximum cluster size was then computed for each iteration and the probability distribution was estimated across the 10,000 iterations. This approach allowed us to determine the minimum cluster size that controls for false positive rate for regions of interest. Parameter estimates were extracted from significant clusters to characterize task-invoked response as a function of WM-load, trait anxiety and groups using 3dmaskave built in AFNI.

Given the prominent effect of long-term stress on drift rate in 2-back condition, we then focused on neural correlates of drift rate in long-term stress and control groups in the following analyses. To identify brain regions showing the interaction effects between group and drift rate, two-sample t-test was conducted for contrast images of the 2-back condition by treating drift rate as a continuous covariate with no mean centering (analogous to ACNOVA). Additionally, we also examined WM-related brain activity associated with drift rate but not necessarily interacting with groups, which might exhibit the mediation effects in the long-term stress different from controls. Whole-brain regression analysis was then conducted to search for the neural correlates of drift rate in the 2-back condition in the stress group. Significant clusters were determined by the same criteria as noted above and corresponding parameter estimates were then extracted. Once associations between drift rate and brain activation were identified, we then conducted correlation analyses to examine whether such brain activation was also associated with individual differences in trait anxiety for both two groups. If WM-related brain activity was correlated with both drift rate and trait anxiety, structural equation models were then constructed to examine the potential mediation effects of WM-related brain activity in these regions.

### Structural equation modeling (SEM)

Separate multi-group SEMs were conducted via MPLUS 7.4 ^65^ to test the mediating effects of WM-related activity (i.e., IPS and MFG) on the association between trait anxiety and drift rate in long-term stress and control groups. Both direct and indirect effects of two groups and their group differences were estimated using 95% bias-corrected CIs with 10000 bootstrapped resamples^66^. The 95% bias-corrected CIs without the inclusion of 0 indicates a statistically significant indirect effect at *P* < 0.05^66^. Several fit indices evaluating the fitness of the proposed models were provided and used the following guidelines for judging good fit: The root mean square error of approximation (RMSEA) is considered adequate below 0.08. The standardized root mean square residual (SRMR) refers to the standardized difference between the observed correlation and the predicted correlation, and considered acceptable with values of 0.08 or less^67^. The comparative fit index (CFI) considers the number of parameters, or paths, in the model and is considered good at 0.93 or above. Parallel analyses were further conducted for state anxiety.

### Prediction analysis

Prediction analyses were performed by Python package “sklearn”, using a machine learning approach with balanced 4-fold cross-validation with 4 repeats combined with linear regression to confirm the conventional correlations^68^. The 4-fold cross-validation procedure was used to avoid overfitting that can occur when the leave-one-out cross-validation procedure is used on small sample sizes. We first estimated r_(predicted, observed)_, the correlation between the values predicted by the regression model and the observed/actual values, using a balanced fourfold cross-validation procedure. The r_(predicted, observed)_ is a measure of how well the independent variable(s) predict the dependent variable. Data were divided into four folds such that the distributions of dependent and independent variables were balanced across folds. A linear regression model was built using three folds, leaving out one fold. The samples in the left-out fold were then predicted using this model, and the predicted values were noted. This procedure was repeated four times, and finally an r_(predicted, observed)_was computed based on the predicted and observed values. Finally, the statistical significance of the model was assessed using nonparametric analysis. The empirical null distribution of r_(predicted, observed)_ was estimated by generating 500 surrogate datasets under the null hypothesis that there was no association between independent and dependent variables. Each surrogate dataset Di of size equal to the observed dataset was generated by permuting the labels (dependent variables) on the observed data points. r_(predicted, observed)i_ was computed using the actual labels of Di and predicted labels using the fourfold cross-validation procedure described previously. This procedure produces a null distribution of r_(predicted, observed)_ for the regression model. The statistical significance of the model was then determined by counting the number of r_(predicted, observed)i_ greater than r_(predicted, observed)_and then dividing that count by the number of Di datasets (500 in our case).

### Network analysis for task-state fMRI data

#### Node definition of brain networks

Core nodes of the typical FPN, DMN and SN were derived from an automated meta-analysis of the most recent 11,406 fMRI studies in Neurosynth (http://www.neurosynth.org)^69^. The nodes in these three networks are presented in **Figure 6a**. Briefly, brain masks of the FPN, DMN and SN were first generated using three separate terms of ‘working memory’, ‘default mode’, and ‘salience network’, respectively. The nodes of the FPN included the DLPFC and the IPS. The nodes in the DMN included the MPFC and the PCC, and the nodes in the SN included the dACC and the AI. Among these ROI masks, the MPFC, PPC and dACC, locating at the middle line structures, form into their own joint clusters across both the left and right hemispheres. For the remaining masks, we combined the clusters from the left and right hemispheres into one unified mask, and time series from the left and right hemispheres were the averaged. The nodes were visualized with the BrainNetViewer (http://www.nitrc.org/projects/bnv/).

#### Intra- and inter-network functional connectivity

Task-specific (i.e., 2-back) ROI-ROI functional connectivity analysis were performed using the CONN toolbox (https://www.nitrc.org/projects/conn/)^70^. Our network coupling metrics derived from the CONN package actually assess functional connectivity between task-invoked time series of certain given regions in each condition separately. This measure is believed to refect functional coupling among brain regions or nodes of interest under certain cognitive task. In this view, we thus feel that the 2-back condition alone rather than the difference between 2- vs. 0-back condition would be better to refect FNP-DMN functional coupling, as the 0-back may not be optimal to serve as a baseline in the context of task-dependent functional connectivity. For each participant, six ROIs’ averaged time series were generated as regressors of interest. Nuisance covariates including cerebrospinal fluid (CSF), white matter (WM) and movement parameters were regressed from the BOLD signal using CompCor method implemented in CONN. Bivariate correlations were then computed between each pair of nodes, resulting in 6 x 6 correlation matrix for each participant in the 0-back and 2-back conditions separately (*Fig. S11*).

Intra- and inter-network functional connectivity metrics were computed separately. For each participant, the intra-network connectivity strength in each condition was calculated by averaging Fisher-z transformed bivariate correlation coefficients between the weighted BOLD time series of nodes within each network. Correspondingly, the inter-network connectivity strength was computed by averaging Fisher-z transformed bivariate correlation coefficients of the weighted BOLD time series across nodes between two different networks. To further investigate how trait anxiety modulates the intra- and inter-network coupling patterns, regression analysis for connectivity and trait anxiety was performed by controlling state anxiety. We finally used regression analysis to explore its relationship with drift rate.

## Author contributions

S.Q. & J.W. designed the research; G.H., J.W., C.L., Y.L., and J.L. performed the research; L.L. analyzed the data; L.L., and S.Q. wrote the manuscript.

## Acknowledgments

This work was supported by the National Natural Science Foundation of China (31522028, 81571056, 2014NT15, 31771246) and the Fundamental Research Funds for the Central Universities. We thank the editor and anonymous reviewers for their valuable comments to improve the manuscript. We also thank Drs. Gang Chen, Baojuan Ye, Ling Wang, Hongyun Liu and Junhao Pan for their advice on modeling and thresholding methods, and Drs. Christina Young and Peter Bayley for language editing.

## Code availability

The scripts for this study are available at https://github.com/QinBrainLab/2018_stress_anxiety_brain.

## Data availability

All the data used in this study are available from the corresponding author upon reasonable request.

## Footnotes

* The final sample size was included for further fMRI data analyses in the stress group.

† The final sample size in the control group. Details are provided in the Methods.

## References

1. de Kloet ER, Joëls M, Holsboer F. Stress and the brain: from adaptation to disease. Nat Rev Neurosci 6, 463–475 (2005).

2. Arnsten AFT. Stress signalling pathways that impair prefrontal cortex structure and function. Nat Rev Neurosci 10, 410–422 (2009).

3. Lupien SJ, McEwen BS, Gunnar MR, Heim C. Effects of stress throughout the lifespan on the brain, behaviour and cognition. Nat Rev Neurosci 10, 434–445 (2009).

4. Joëls M, Pu Z, Wiegert O, Oitzl MS, Krugers HJ. Learning under stress: how does it work? Trends in cognitive sciences 10, 152–158 (2006).

5. McEwen BS, Morrison JH. The brain on stress: vulnerability and plasticity of the prefrontal cortex over the life course. Neuron 79, 16–29 (2013).

6. Franklin TB, Saab BJ, Mansuy IM. Neural mechanisms of stress resilience and vulnerability. Neuron 75, 747–761 (2012).

7. Weger M, Sandi C. High anxiety trait: a vulnerable phenotype for stress-induced depression. Neuroscience & Biobehavioral Reviews 87, 27–37 (2018).

8. Bishop S, Forster S. Trait anxiety, neuroticism, and the brain basis of vulnerability to affective disorder. In: The Cambridge handbook of human affective neuroscience. (ed^(eds). Cambridge University Press (2013).

9. Edwards EJ, Edwards MS, Lyvers M. Cognitive trait anxiety, situational stress, and mental effort predict shifting efficiency: Implications for attentional control theory. Emotion 15, 350 (2015).

10. Calvo MG, Eysenck MW, Ramos PM, Jiménez A. Compensatory reading strategies in test anxiety. Anxiety, Stress, and Coping 7, 99–116 (1994).

11. Calvo MG. Phonological working memory and reading in test anxiety. Memory 4, 289–306 (1996).

12. Bishop SJ. Neurocognitive mechanisms of anxiety: an integrative account. Trends in Cognitive Sciences 11, 307–316 (2007).

13. Browning M, Behrens TE, Jocham G, O’Reilly JX, Bishop SJ. Anxious individuals have difficulty learning the causal statistics of aversive environments. Nat Neurosci 18, 590–596 (2015).

14. Ratcliff R. A theory of memory retrieval. Psychol Rev 85, 59–108 (1978).

15. Ratcliff R, McKoon G. The diffusion decision model: theory and data for two-choice decision tasks. Neural computation 20, 873–922 (2008).

16. Mulder MJ, van Maanen L, Forstmann BU. Perceptual decision neurosciences —— A model-based review. Neuroscience 277, 872–884 (2014).

17. Sereno M, Pitzalis S, Martinez A. Mapping of contralateral space in retinotopic coordinates by a parietal cortical area in humans. Science 294, 1350–1354 (2001).

18. Liu T, Pleskac TJ. Neural correlates of evidence accumulation in a perceptual decision task. Journal of neurophysiology 106, 2383–2398 (2011).

19. Roitman JD, Shadlen MN. Response of Neurons in the Lateral Intraparietal Area during a Combined Visual Discrimination Reaction Time Task. The Journal of Neuroscience 22, 9475 (2002).

20. Birnbaum SG, et al. Protein Kinase C Overactivity Impairs Prefrontal Cortical Regulation of Working Memory. Science 306, 882 (2004).

21. Wang M, et al. α2A-Adrenoceptors Strengthen Working Memory Networks by Inhibiting cAMP-HCN α Channel Signaling in Prefrontal Cortex. Cell 129, 397–410 (2007).

22. Bishop SJ. Trait anxiety and impoverished prefrontal control of attention. Nat Neurosci 12, 92–98 (2009).

23. Raichle ME, MacLeod AM, Snyder AZ, Powers WJ, Gusnard DA, Shulman GL. A default mode of brain function. Proceedings of the National Academy of Sciences 98, 676 (2001).

24. Anticevic A, Cole MW, Murray JD, Corlett PR, Wang X-J, Krystal JH. The role of default network deactivation in cognition and disease. Trends in cognitive sciences 16, 584–592 (2012).

25. Menon V. Large-scale brain networks and psychopathology: a unifying triple network model.

26. Luo Y, Qin S, Fernández G, Zhang Y, Klumpers F, Li H. Emotion perception and executive control interact in the salience network during emotionally charged working memory processing. Hum Brain Mapp 35, 5606–5616 (2014).

27. Qin S, Hermans EJ, van Marle HJF, Luo J, Fernández Gn. Acute Psychological Stress Reduces Working Memory-Related Activity in the Dorsolateral Prefrontal Cortex. Biol Psychiat 66, 25–32 (2009).

28. Cocchi L, Zalesky A, Fornito A, Mattingley JB. Dynamic cooperation and competition between brain systems during cognitive control. Trends in Cognitive Sciences 17, 493–501 (2013).

29. Sridharan D, Levitin DJ, Menon V. A critical role for the right fronto-insular cortex in switching between central-executive and default-mode networks. Proceedings of the National Academy of Sciences 105, 12569 (2008).

30. Hermans EJ, Henckens MJAG, Joëls M, Fernández Gn. Dynamic adaptation of large-scale brain networks in response to acute stressors. Trends in Neurosciences 37, 304–314 (2014).

31. Duan H, et al. Chronic stress exposure decreases the cortisol awakening response in healthy young men. Stress 16, 630–637 (2013).

32. Duan H, Yuan Y, Yang C, Zhang L, Zhang K, Wu J. Anticipatory processes under academic stress: An ERP study. Brain Cognition 94, 60–67 (2015).

33. Assessment of state and trait anxiety: Conceptual and methodological issues. Louisiana Psychological Association (1985).

34. Hildenbrand AK, Nicholls EG, Aggarwal R, Brody-Bizar E, Daly BP. Symptom Checklist-90-Revised (SCL-90-R). The Encyclopedia of Clinical Psychology, 1–5 (2015).

35. MacLeod C, Hagan R. Individual differences in the selective processing of threatening information, and emotional responses to a stressful life event. Behav Res Ther 30, 151–161 (1992).

36. Derakshan N, Eysenck MW. Anxiety, processing efficiency, and cognitive performance: New developments from attentional control theory. European Psychologist 14, 168–176 (2009).

37. Qin S, et al. The effect of moderate acute psychological stress on working memory-related neural activity is modulated by a genetic variation in catecholaminergic function in humans. Frontiers in Integrative Neuroscience 6, 16 (2012).

38. Schoofs D, Pabst S, Brand M, Wolf OT. Working memory is differentially affected by stress in men and women. Behavioural Brain Research 241, 144–153 (2013).

39. Quaedflieg CW, Schwabe L. Memory dynamics under stress. Memory 26, 364–376 (2018).

40. Ebner K, Singewald N. Individual differences in stress susceptibility and stress inhibitory mechanisms. Curr Opin Behav Sci 14, 54–64 (2017).

41. Eysenck MW, Derakshan N, Santos R, Calvo MG. Anxiety and cognitive performance: attentional control theory. Emotion 7, 336 (2007).

42. Phillips ML, Ladouceur CD, Drevets WC. A neural model of voluntary and automatic emotion regulation: implications for understanding the pathophysiology and neurodevelopment of bipolar disorder. Mol Psychiatr 13, 833–857 (2008).

43. Goldin PR, McRae K, Ramel W, Gross JJ. The Neural Bases of Emotion Regulation: Reappraisal and Suppression of Negative Emotion. Biol Psychiat 63, 577–586 (2008).

44. Menon V, Uddin LQ. Saliency, switching, attention and control: a network model of insula function. Brain Structure and Function 214, 655–667 (2010).

45. Pessoa L. How do emotion and motivation direct executive control? Trends in cognitive sciences 13, 160–166 (2009).

46. Whitfield-Gabrieli S, Ford JM. Default mode network activity and connectivity in psychopathology. Annual review of clinical psychology 8, 49–76 (2012).

47. Roitman JD, Shadlen MN. Response of neurons in the lateral intraparietal area during a combined visual discrimination reaction time task. J Neurosci 22, 9475–9489 (2002).

48. Ratcliff R, Smith PL. A comparison of sequential sampling models for two-choice reaction time. Psychol Rev 111, 333 (2004).

49. Rasmussen K, Morilak DA, Jacobs BL. Single unit activity of locus coeruleus neurons in the freely moving cat: I. During naturalistic behaviors and in response to simple and complex stimuli. Brain Research 371, 324–334 (1986).

50. Hampson M, Driesen N, Roth JK, Gore JC, Constable RT. Functional connectivity between task-positive and task-negative brain areas and its relation to working memory performance. Magnetic resonance imaging 28, 1051–1057 (2010).

51. Chen AC, et al. Causal interactions between fronto-parietal central executive and default-mode networks in humans. Proceedings of the National Academy of Sciences 110, 19944 (2013).

52. Etkin A, Wager TD. Functional Neuroimaging of Anxiety: A Meta-Analysis of Emotional Processing in PTSD, Social Anxiety Disorder, and Specific Phobia. Am J Psychiat 164, 1476–1488 (2007).

53. Lerche V, Voss A, Nagler M. How many trials are required for parameter estimation in diffusion modeling? A comparison of different optimization criteria. Behavior Research Methods 49, 513–537 (2017).

54. Cavanagh JF, Wiecki TV, Kochar A, Frank MJ. Eye tracking and pupillometry are indicators of dissociable latent decision processes. Journal of Experimental Psychology: General 143, 1476–1488 (2014).

55. O’Callaghan C, et al. Visual Hallucinations Are Characterized by Impaired Sensory Evidence Accumulation: Insights From Hierarchical Drift Diffusion Modeling in Parkinson ÇÔs Disease.Biological Psychiatry: Cognitive Neuroscience and Neuroimaging 2, 680–688 (2017).

56. Liston C, McEwen BS, Casey BJ. Psychosocial stress reversibly disrupts prefrontal processing and attentional control. Proceedings of the National Academy of Sciences 106, 912 (2009).

57. Derogatis LR, Unger R. Symptom Checklist-90-Revised. The Corsini Encyclopedia of Psychology, 1–2 (2010).

58. Wiecki T, Sofer I, Frank M. HDDM: Hierarchical Bayesian estimation of the Drift-Diffusion Model in Python. Front Neuroinform 7, 14 (2013).

59. Ratcliff R, McKoon G. The Diffusion Decision Model: Theory and Data for Two-Choice Decision Tasks. Neural Computation 20, 873–922 (2007).

60. Ratcliff R, Huang-Pollock C, McKoon G. Modeling individual differences in the go/no-go task with a diffusion model. Decision 5, 42–62 (2018).

61. Zhang J, et al. Different decision deficits impair response inhibition in progressive supranuclear palsy and Parkinson ÇÔs disease. Brain 139, 161–173 (2015).

62. Zhang J, Rowe JB. Dissociable mechanisms of speed-accuracy tradeoff during visual perceptual learning are revealed by a hierarchical drift-diffusion model. Front Neurosci-Switz 8, 69 (2014).

63. Singmann H, et al. afex: Analysis of factorial experiments. R package version 0.16-1. R Package Version 016 1, (2016).

64. Nichols T, Brett M, Andersson J, Wager T, Poline J-B. Valid conjunction inference with the minimum statistic. Neuroimage 25, 653–660 (2005).

65. Hayes AF, Preacher KJ, Myers TA. Mediation and the estimation of indirect effects in political communication research. Sourcebook for political communication research: Methods, measures, and analytical techniques 23, 434–465 (2011).

66. Preacher KJ, Hayes AF. Asymptotic and resampling strategies for assessing and comparing indirect effects in multiple mediator models. Behavior research methods 40, 879–891 (2008).

67. Hu L, Bentler PM. Cutoff criteria for fit indexes in covariance structure analysis: Conventional criteria versus new alternatives. Structural Equation Modeling: A Multidisciplinary Journal 6, 1–55 (1999).

68. Cohen JR. Decoding developmental differences and individual variability in response inhibition through predictive analyses across individuals. Front Hum Neurosci 4, 47 (2010).

69. Yarkoni T, Poldrack RA, Nichols TE, Van Essen DC, Wager TD. Large-scale automated synthesis of human functional neuroimaging data. Nature Methods 8, 665–670 (2011).

70. Whitfield-Gabrieli S, Nieto-Castanon A. Conn: A Functional Connectivity Toolbox for Correlated and Anticorrelated Brain Networks. Brain Connectivity 2, 125–141 (2012).

