## Supplementary Materials for "Long-term stress and trait anxiety alter brain network balance in dynamic decisions during working memory"

**Supplemental Materials**

**Supplementary Texts**

**Bayesian hierarchical version of Drift Diffusion Model (HDDM)**

We applied a Bayesian hierarchical version of Drift Diffusion Model (HDDM) to decompose our observed choice responses (i.e., RTs) into latent dynamic computation processes modulated by four free parameters: drift rate *v*, decision threshold *a*, starting point *z*, and non-decision time *t*. An analytic solution to the probability distribution of the termination times (i.e., RTs) was provided by the following formula (Wiener diffusion model):

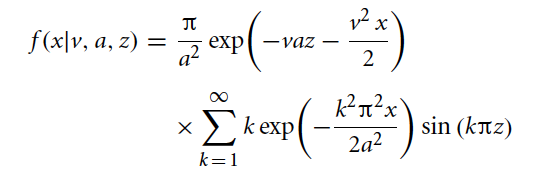

Where *x* represents a certain RT for a given participant, *v* represents drift rate, *a* represents decision boundary, and *z* represents the prior response bias. It is worth to note that this is a probability density function, which can be interpreted as follows: For each participant under certain condition (e.g., 2-back), given *v*, *a* and *z*, what’s the probability that the real RT equals to *x*? Later on, the drift diffusion model (DDM) has been extended to include additional noise parameters that can capture inter-trial variability in drift rate, non-decision time and the starting point in order to account for two phenomena observed in decision making tasks: most notably cases where errors are faster or slower than correct responses. Models that take this critical feature into account are referred to as the full DDM ^1^. The equation in the primitive version of the DDM above for the probability distribution of the termination times assume that time begins at zero — that is, non-decision time *t* equals zero. To introduce a non-zero value for non-decision time *t* into the equation, each occurrence of *x* would be replaced by *x – t*. Then, *x* would refer to reaction times with a non-decision component equal to *t*. This produces a full DDM: f(x, v ,a ,z ,t), which is used for modeling. Since the formula contains an infinite sum, the HDDM uses a likelihood function as formulated by *Navarro and Fuss (2009)^2^*. We here denote this likelihood function as $f|\theta$ for the illustrative purpose, which represents the likelihood of observing the data (in this case RTs) given each parameter value. In other words, observed data points of each participant x_i,j_ (where i = 1, …, S_j_ data points per participant, and j = 1, …, N for N participants) are distributed according to the likelihood function $f|\theta$. As noted above, parameters from individual participants are not completely independent, but drawn from the group distribution. Each participant’s parameters θ_j_ are assumed to obey a normal distribution around a group mean with a specific group variance [λ = (μ,σ), where these group parameters are estimated from the data given hyper-priors G0], resulting in the following generative description:

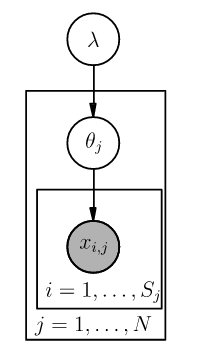
μ,σ ∼ G_0_()

θj ∼ N(μ, σ^2^)

x_i,j_ ∼ f(θj)

Circles represent continuous random variables. Arrows connecting circles specify conditional dependence between random variables. Shaded circles represent the observed data. The plates around graphical nodes mean that multiple identical, independent distributed random variables exist. From this graphical notation, we can clearly see that the Bayesian method implemented in the HDDM lends itself naturally to a hierarchical design, allowing group and subject parameters to be estimated simultaneously at different hierarchical levels. The P(x|θ) can be calculated from the likelihood function mentioned above, by specifying the prior beliefs $P(\theta)$^3^, we can use Bayes formula to make inference on the probability of parameters θ: P(θ|x) = $\frac{P\left( x | \theta\right) P(\theta)}{P(x)}$. Finally, each participant obtained four parameters estimated from above Hierarchical Bayesian Estimation process under certain condition.

Unlike an unbiased response bias across conditions and groups used in previous studies (i.e., fixed to 0.5) ^4, 5, 6^, we here assumed that the response bias would not vary as a function of conditions, but it would vary across groups only. For instance, participants would not bias toward pressing the button in the 0-back condition or withholding response in the 2-back condition, but participants under long-term stress may behave differently from the controls. Indeed, model comparisons provided the best fit and good convergence for the model allowing for changes in model parameters including drift rate *v*, decision threshold *a*, non-decision time *t* between conditions and groups. And response bias *z* varied across groups rather than conditions (**Fig.S4 & Table S4**). For data in **Table S4,** we thus conducted 2 (Group)-by-2 (WM-load) ANOVA for drift rate *v*, decision threshold *a*, and non-decision time *t* to examine the main effects of long-term stress and WM loads, and their interaction effects. For the response bias *z*, however, we examined the group difference between long-term stress and controls by employing independent two-sample t test.

**Supplementary Results**

**Correlations of WM performance with computational, neural activation and connectivity**

To examine the relations of latent-, neural-, and connectivity-measures with actual behavioural performance (i.e., RTs and accuracy), we conducted a set of exploratory correlation analyses for these variables. Given that we observed prominent long-term stress effect on averaged RTs in the 2-back condition, we thus restricted our analyses on data in the condition. As shown in **Table S5**, these analyses revealed significantly negative correlations between RTs and drift rate (v) in both the stress and control groups under the 2-back condition [stress: r(34) = -0.43, p = 0.008; control: r(30) = -0.68, p < 0.001]*.* The averaged RTs also exhibited a positive correlation with decision-threshold (a) [stress: r(34) = 0.94, p < 0.001; control: r(30) = 0.81, p < 0.001]*.* We did not observe any reliable correlations of averaged RTs with task-induced activity in the anterior insula showing the main effect of long-term stress and brain regions of the DMN and SN showing the Group-by-WM interaction effects. No correlations were found in the IPS and MFG with RTs, nor SN-DMN coupling and FPN-DMN decoupling with RTs. These results indicate that the long-term stress effect on RTs in the 2-back condition was closely linked to drift rate and decision-threshold, with higher drift rate and lower decision-threshold associated with faster RT.

For averaged accuracy measurement, we found that individuals with higher accuracy under long-term stress was associated with less activity in the right anterior insula [stress: r(34) = -0.48, p = 0.003; control: r(30) = 0.15, p = 0.42; a group difference: z = -2.65, p = 0.008] and weaker SN-DMN coupling that also exhibited the main effect of long-term stress [SN-DMN coupling: stress: r(34) = -0.49, p = 0.002; control: r(30) = 0.15, p = 0.42; a group difference: z = -2.7, p = 0.007]. Likewise, less MFG activity under long-term stress led to higher level of accuracy [stress: r(34) = -0.37, p = 0.03; control: r(30) = 0.06, p = 0.74; no group difference: z = -1.76, p = 0.08]. Controls with higher accuracy exhibited stronger activation in the mPFC of the DMN [stress: r(34) = -0.18, p = 0.29; control: r(30) = 0.36, p = 0.04; a group difference: z = -2.2, p = 0.03]. No other brain regions in the DMN and SN were associated with accuracy, and no evidence shows any correlations of accuracy with the IPS activity and FPN-DMN decoupling. Latent measures such as drift rate and decision threshold exhibited no correlation with accuracy (*see* ***Table S9***). Together, individuals with higher accuracy were associated with less activity in the right anterior insula and weaker SN-DMN coupling under long-term stress but not controls. These observations appear consistent with our speculation of enhanced processing efficiency under long-term stress.

**Correlations of drift rate and psychological distress with neural activation during WM under long-term stress**

To examine whether neural activation during WM corrrelated with drift rate under long-term stress, we conducted two additional analyses at both regions of interest (ROIs) and the whole-brain levels. For the ROI level, we computed the Pearson’s correlations between drift rate and hyper-activation in the insula in the 0- and 2-back conditions in both stress and control groups. As shown in **Table S14** below, we did not observe any reliable correlations. On the whole-brain level, we conducted separate regression analyses for brain activity maps in 0-, 2- and 2- vs. 0-back conditions separately. Again, we did not observe any reliable effects in the insula in relation to drift rate using a height threshold of p < 0.005 and a spatial extent threshold of p < 0.05 corrected for these regions of interest (see **Table S15&16**).

To examine whether neural activation during WM corrrelated with psychological distress under long-term stress, we also conducted two additional analyses at both regions of interest (ROIs) and the whole-brain levels. At the ROI level, as shown **Table S17** below we did not observe any reliable correlation between activation in the SN ROIs and psychological distress levels in each group under either condition. At the whole brain level, we conducted separate simple regression analyses for brain activity maps in the contrast of 0-, 2- and 2- vs. 0-back conditions with psychological distress as a covariate of interest. Thse analyses revealed significant clusters in a set of distributedbrain regions of the prefrontal and temporal lobes (**Table S18&19**). But we observed no reliable effect in the typical SN regions including the anterior insula and dACC in relation to psychological distress using the same thresholding criteria as our original manuscript.

**Supplementary Figures S1-S11**

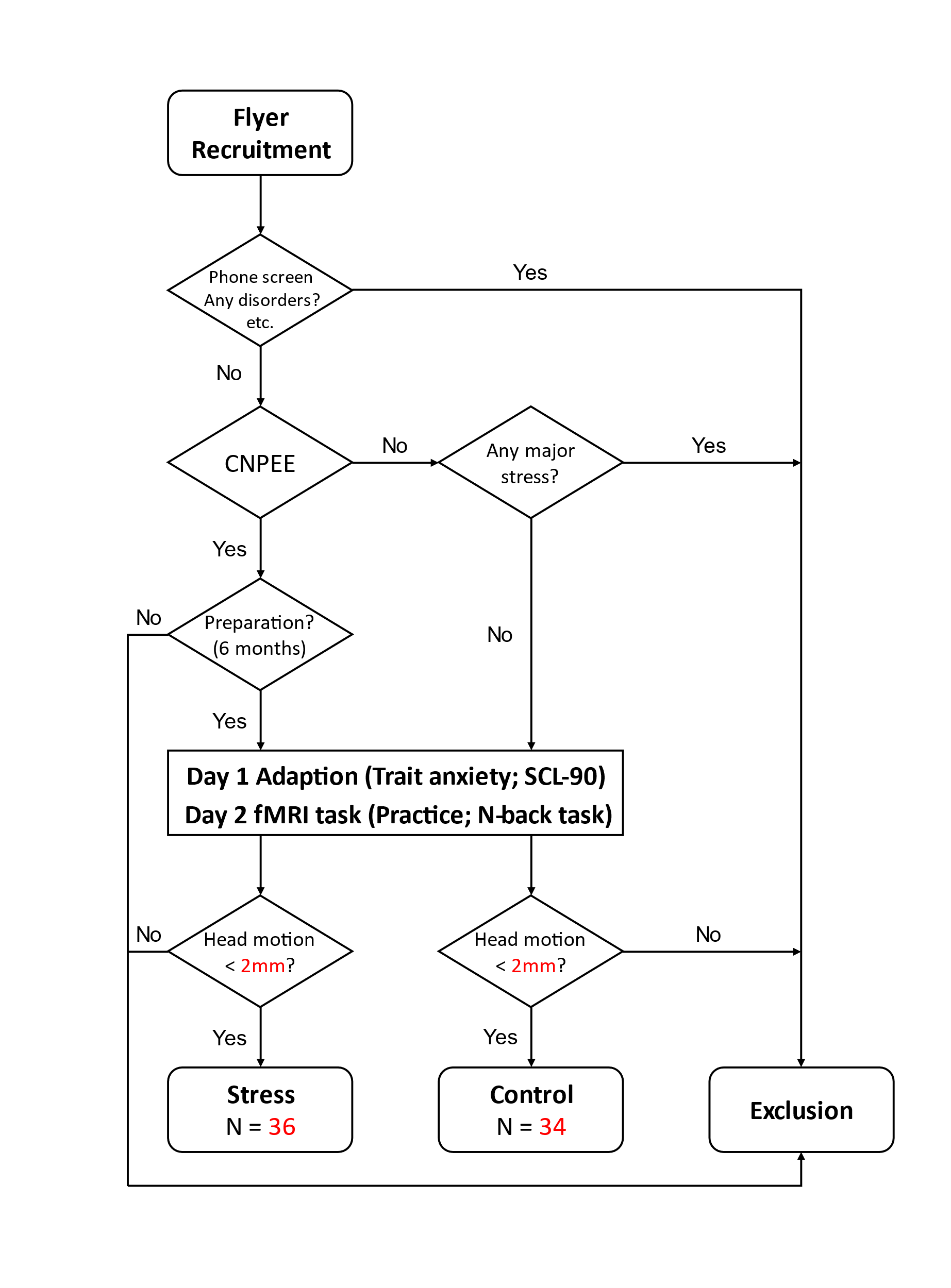
**Figure S1. The chart flow of participant recruitment in our present study.** The detail procedure was described in the Methods part of the manuscript. It is worth to note that all participants were invited to visit the laboratory and fill in anxiety and SCL questionnaires one day (Day 1) prior to the formal fMRI experiment (Day 2). This procedure may allow us to reduce potential confounds by acute stress induced by being fMRI scanning.

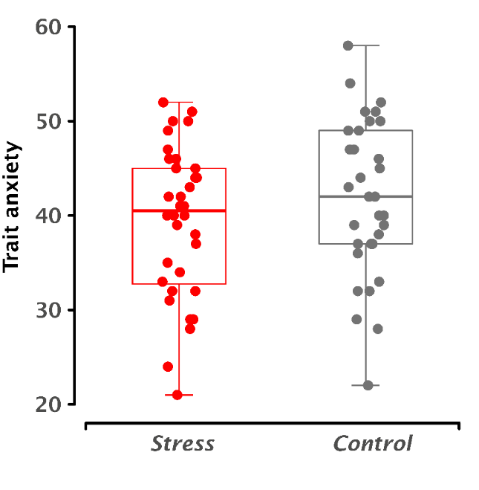

**Figure S2. Boxplot of trait anxiety scores for long-term stress and control groups**. The box plots consist of the minimum, the maximum, the sample median, and the first and third quartiles of trait anxiety. The y-axis represents trait anxiety scores.

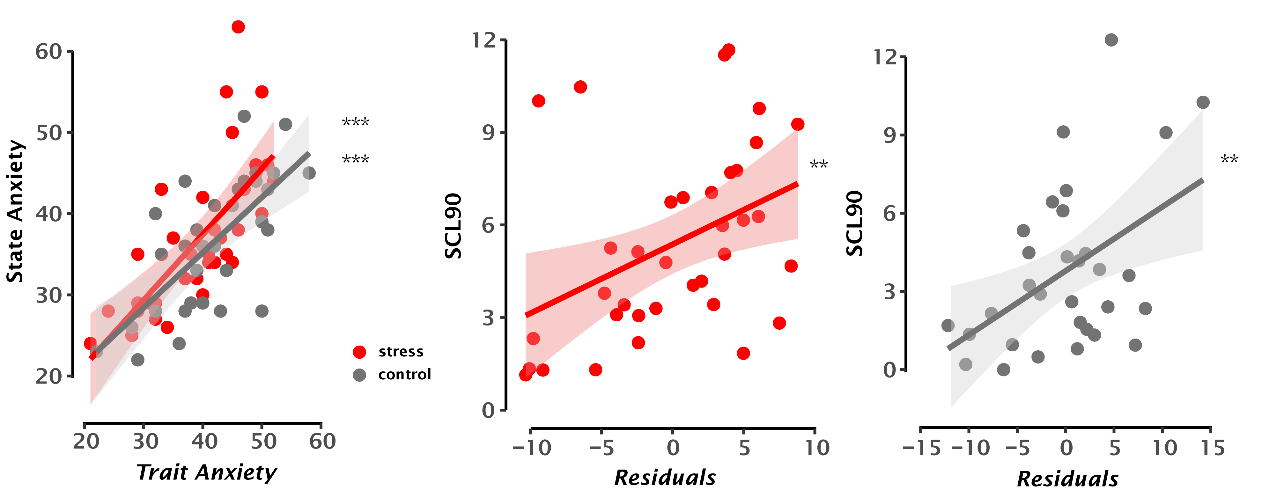

**Figure S3. Correlations between trait anxiety and state anxiety in the long-term stress and control groups (left panel), as well as partial correlations between trait anxiety and psychological distress after regressing out state anxiety in the long-term stress (middle panel) and control groups (right panel).** Scatter plots show the corresponding correlations with the lines of best fit. The shadowed areas represent 95% confidence intervals. The x-axes in the middle and right panels represent residuals of trait anxiety after regressing out state anxiety. Notes: ***P* < 0.005, ****P* < 0.001.

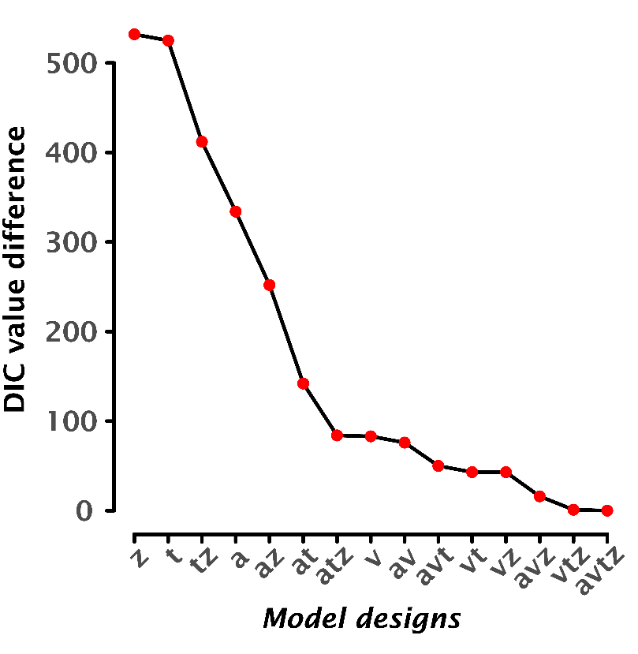

**Figure S4. Model comparisons among 15 model variants.** Model comparisons were conducted by using Deviance Information Criterion (DIC). The y-axis represents the DIC value differences between each of the other model variants and the best fit model. The x-axis represents the number of models with corresponding free parameters. A total of 15 model variants with different parameter constraints were established for participants across both stress and control groups (details provided in **Methods**).

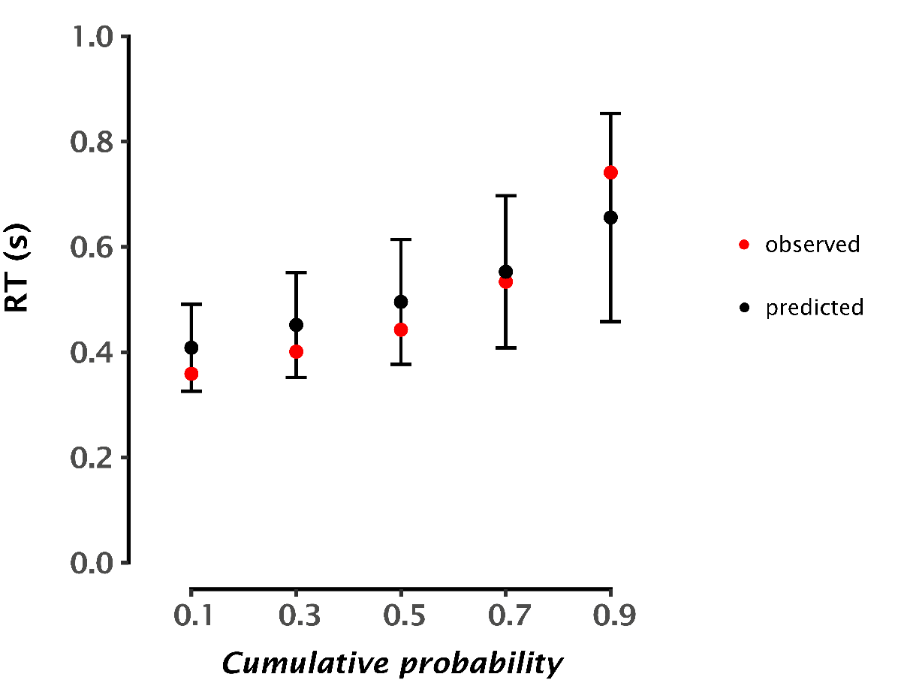

**Figure S5. The quantile probability for posterior predictive checks.** The observed (in red) and predicted (in black) RTs as a function of the cumulative probabilities varying from 10, 30, 50, 70 to 90 percentiles. The quantile function takes the specified probability as input and returns the value X, such that *P(X) = P – value.* The observed data is a fixed value without variance. Error bar represents the standard deviation (SD) of the posterior predictive distribution from the model, which captures the degree to estimation uncertainty.

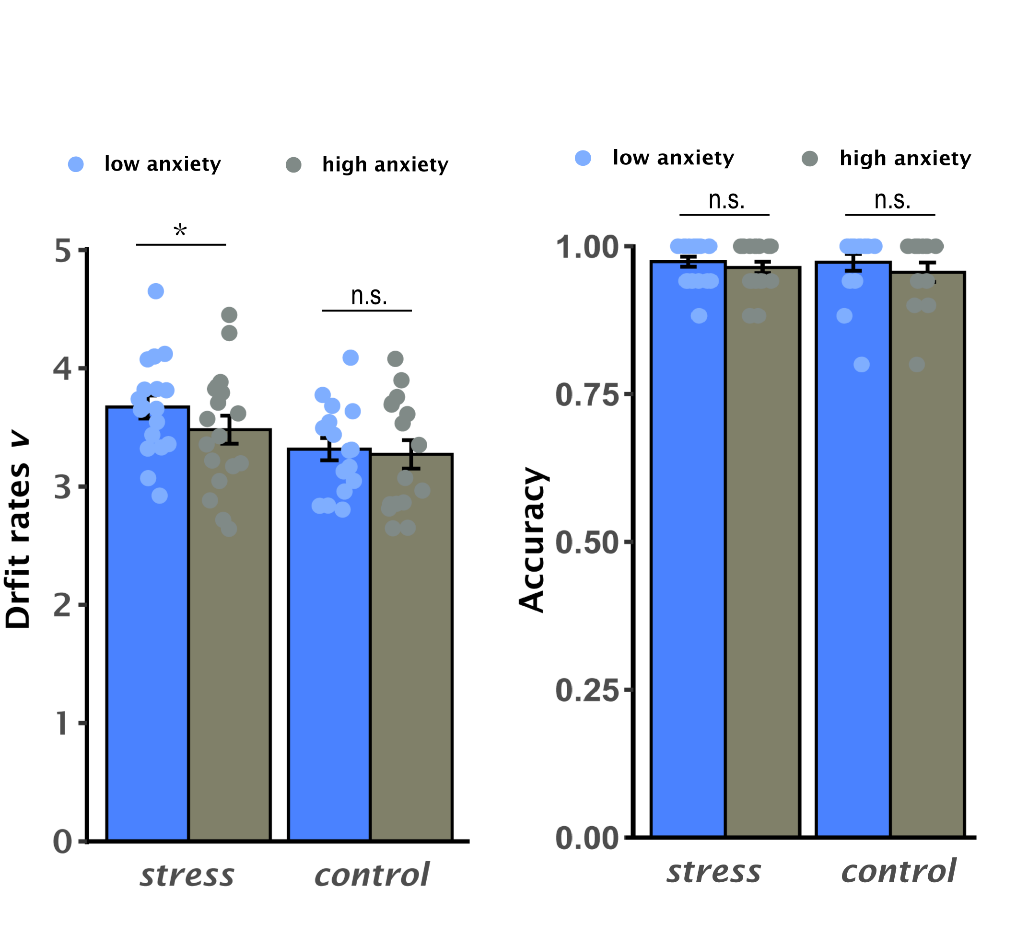

**Figure S6**. **Averaged latent drift rate (left panel) and accuracy (right panel) in the 2-back condition when splitting participants into low and high trait anxiety in both stress and control groups.** This median-split analysis here was conducted for illustration purpose only. Error bar represents standard error of mean. Notes: * p < 0.05; n.s., no significance.

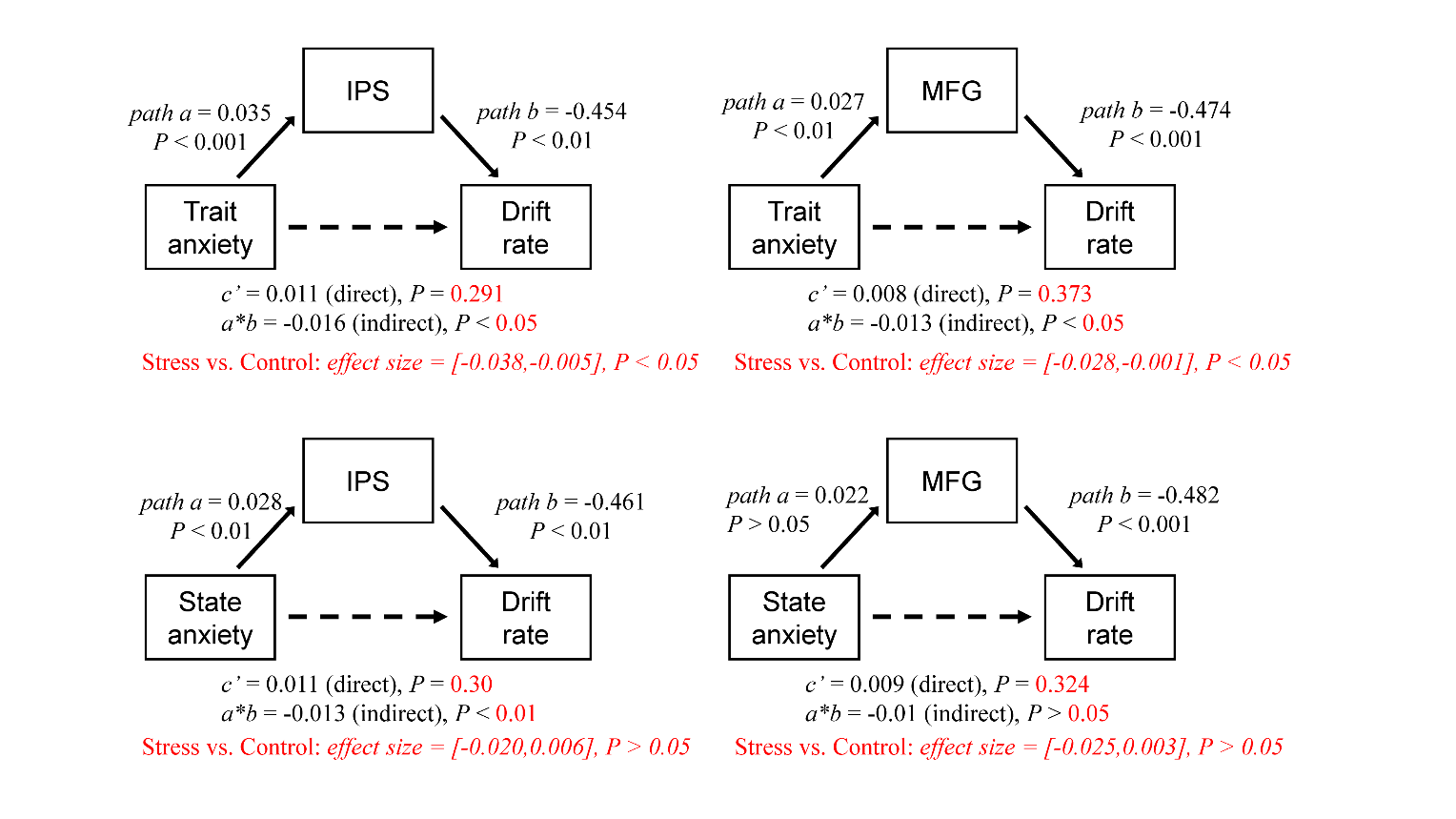

**Figure S7. The mediation effects of WM-related activity in the IPS and MFG on the indirect association between trait (state) anxiety and drift rate in the long-term stress and control groups.** The group differences in the mediation effects were examined using a nonparametric bootstrap resampling approach. The indirect effect was considered significant if the 95% CI did not include zero.IPS, intraparietal sulcus; MFG, middle frontal gyrus; c’, direct path; a*b, indirect path. More details on the statistics are provided in Table S7-8 below.

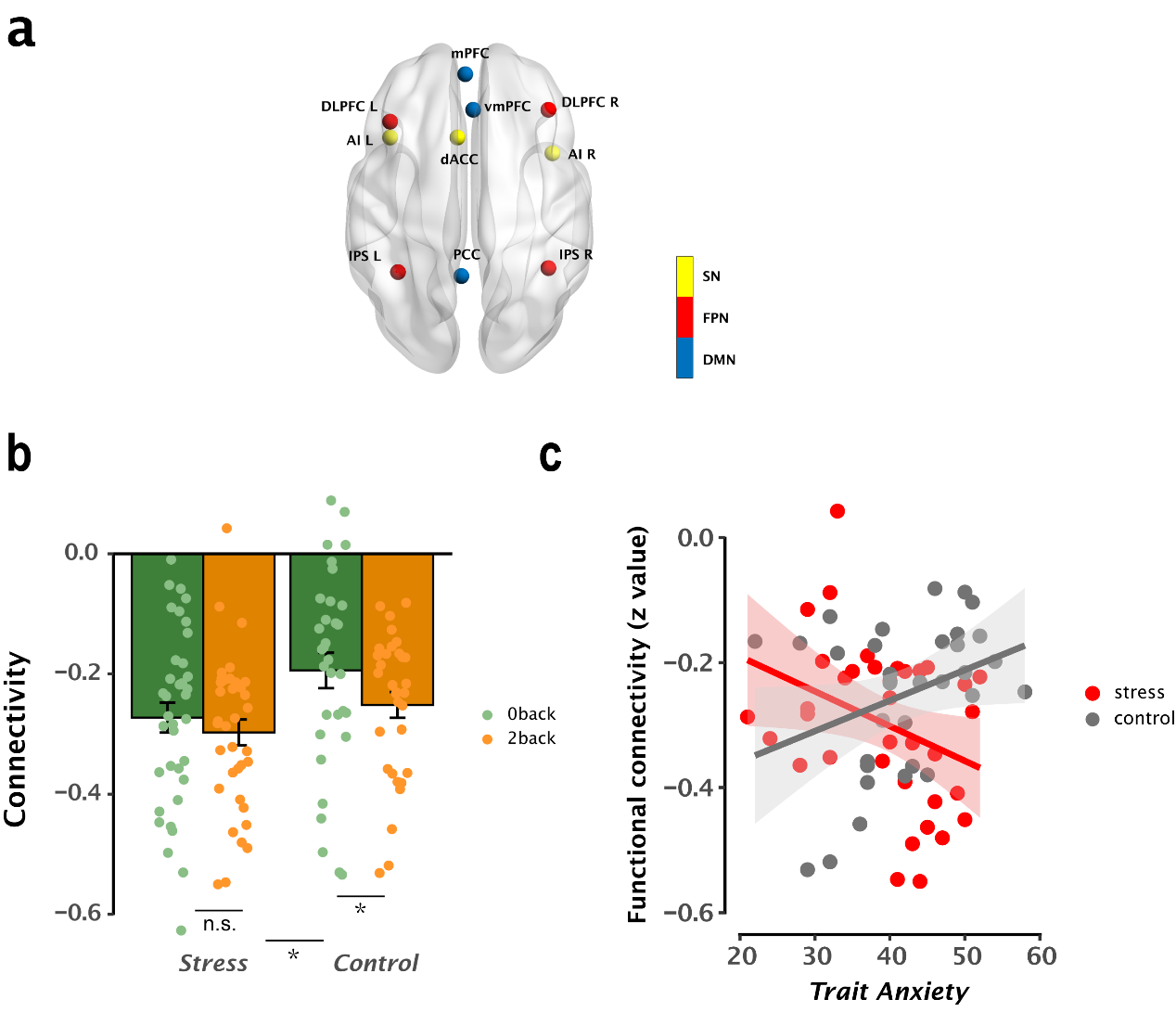

**Figure S8. Inter-network connectivity modulated by trait anxiety under long-term stress.** (**a**) Nodes of the three brain networks involved in WM processing, including the default-node network (DMN, coded in blue), frontal-parietal network (FPN, coded in red) and salience network (SN, coded in yellow). ROIs were defined as spheres with 6-mm around these MNI coordinates (see **Table S20**) derived from previous studies^7, 8, 9^. (**b**) Bar graphs represent FPN-DMN decoupling as a function of 0-and 2-back conditions in the stress and control groups. A 2-by-2 ANOVA revealed a significant main effect of Group, with greater FPN-DMN decoupling under long-term stress than controls [*F(1, 66) = 4.38, p = 0.04*]. Dots represent each participant’s data. (**c**) Scatter plots depict the correlation between trait anxiety and FPN-DMN coupling in the long-term stress and control groups in 2-back condition. The FPN-DMN decoupling was significantly correlated with trait anxiety under long-term stress [r(34) = -0.34, p = 0.04]. Further test for Fisher’s z transformed correlations revealed a significant difference between two groups, indicating an interaction of trait anxiety and long-term stress [z = -2.77, p = 0.006].

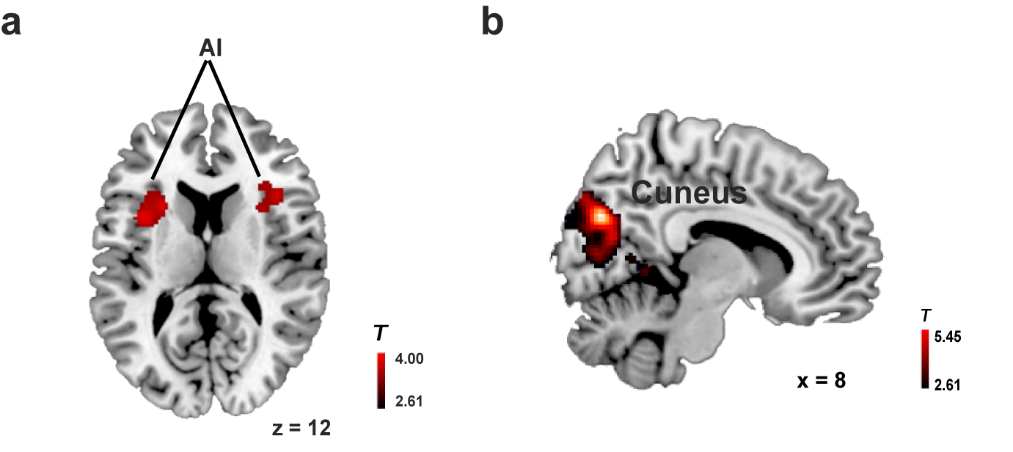

**Figure S9. Brain regions showing the main effect of long-term stress.** Significant clusters in (a) the bilateral anterior insula and (b) the cuneus was derived from the contrast of long-term stress with control group from whole-brain ANOVA with group as between-subject factor (stress vs. control) and WM loads (0- vs. 2-back) as within-subject factor. Both the bilateral anterior insula and the cuneus exhibit the main effect of long-term stress, with general hyper-activation in these regions under long-term stress than controls. Significant clusters were determined by a height threshold p < 0.001 and an extent threshold p < 0.05 corrected. Color bar indicates minimum and maximal T values.

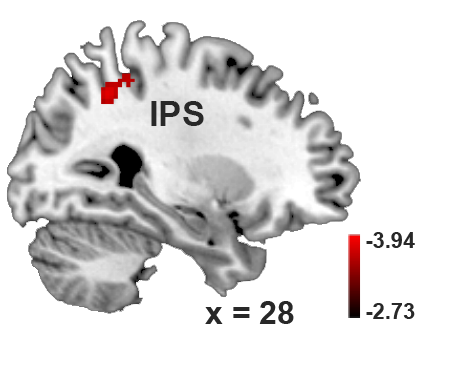

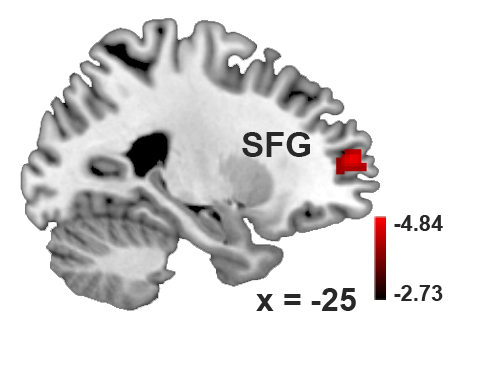

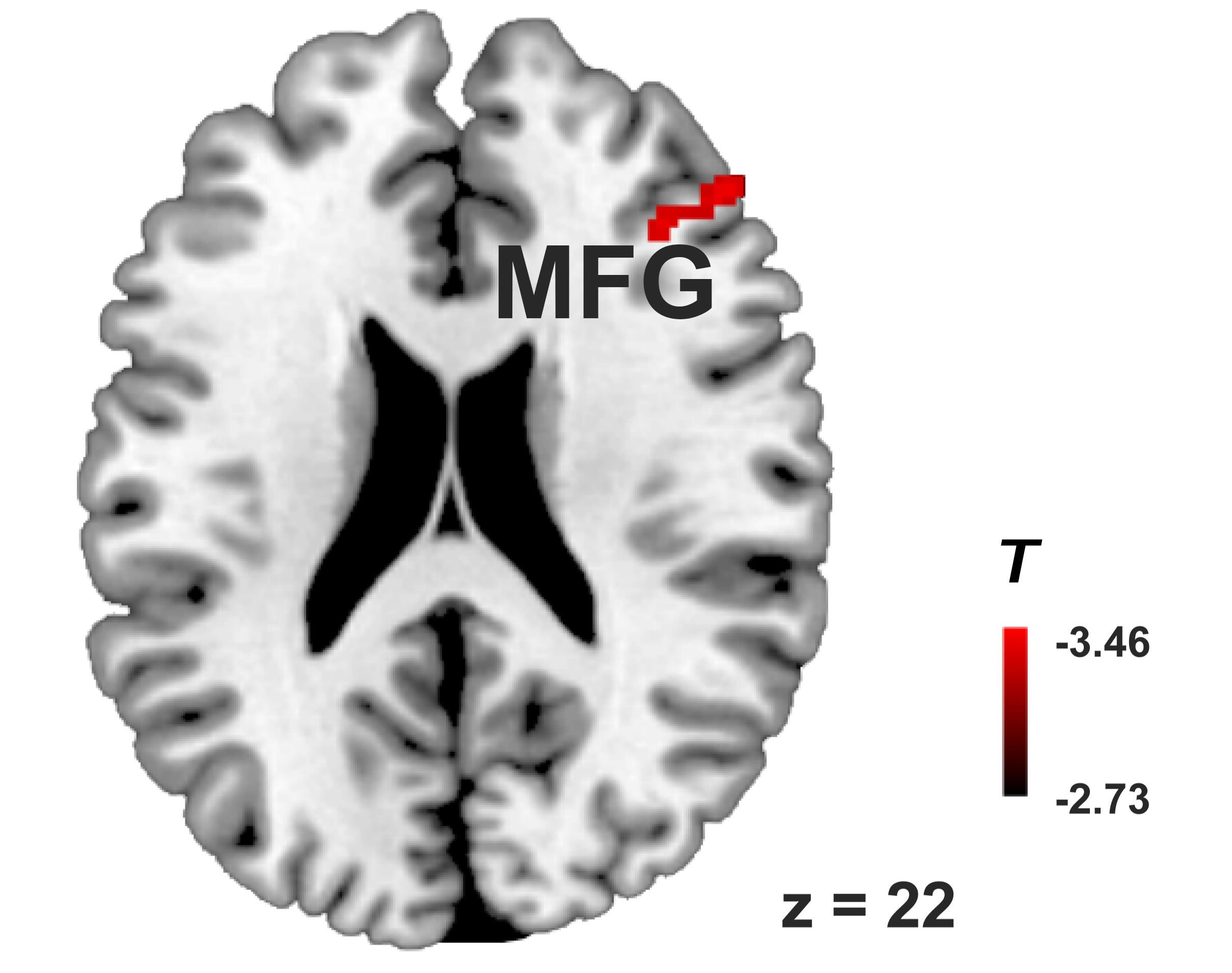

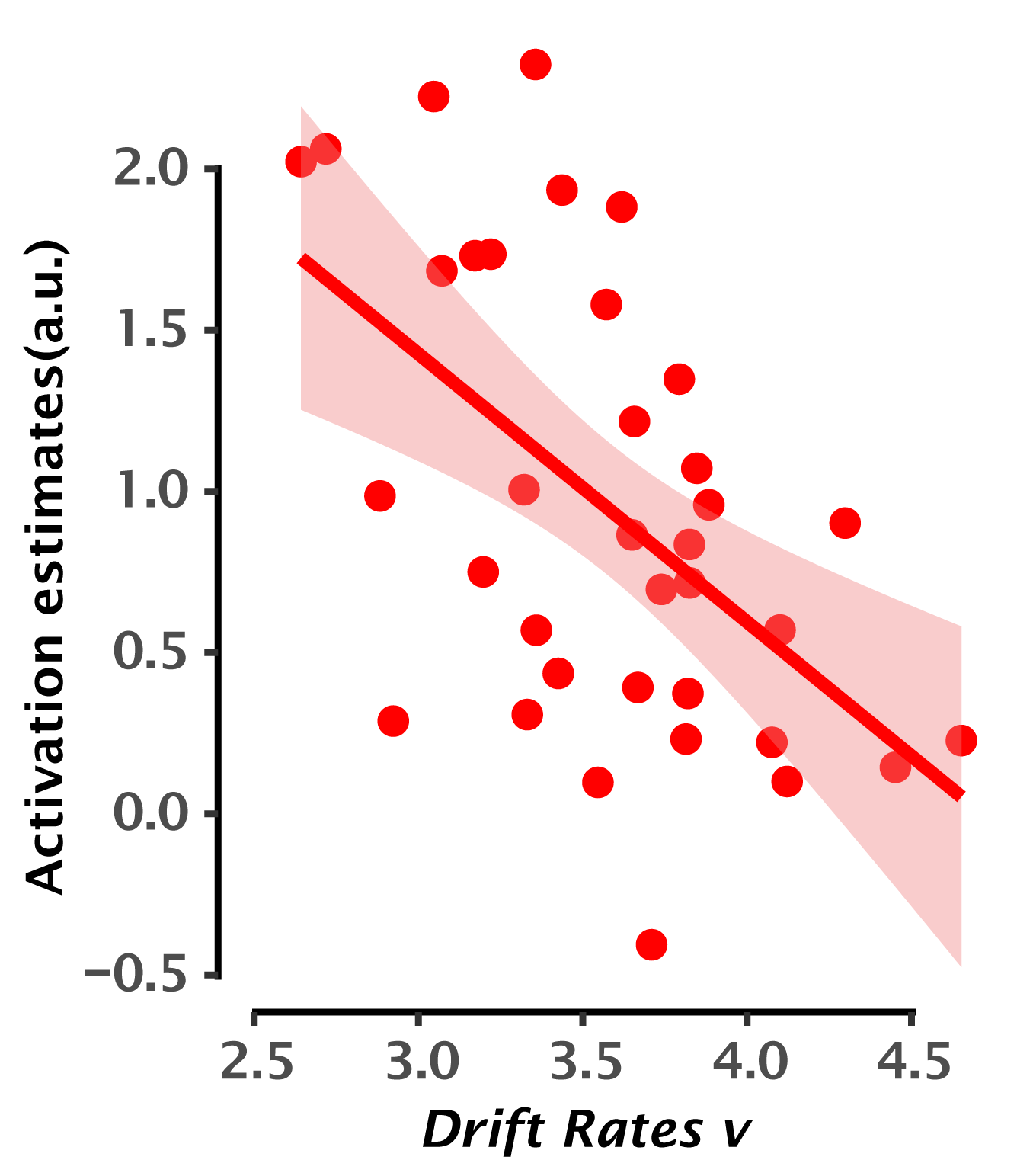

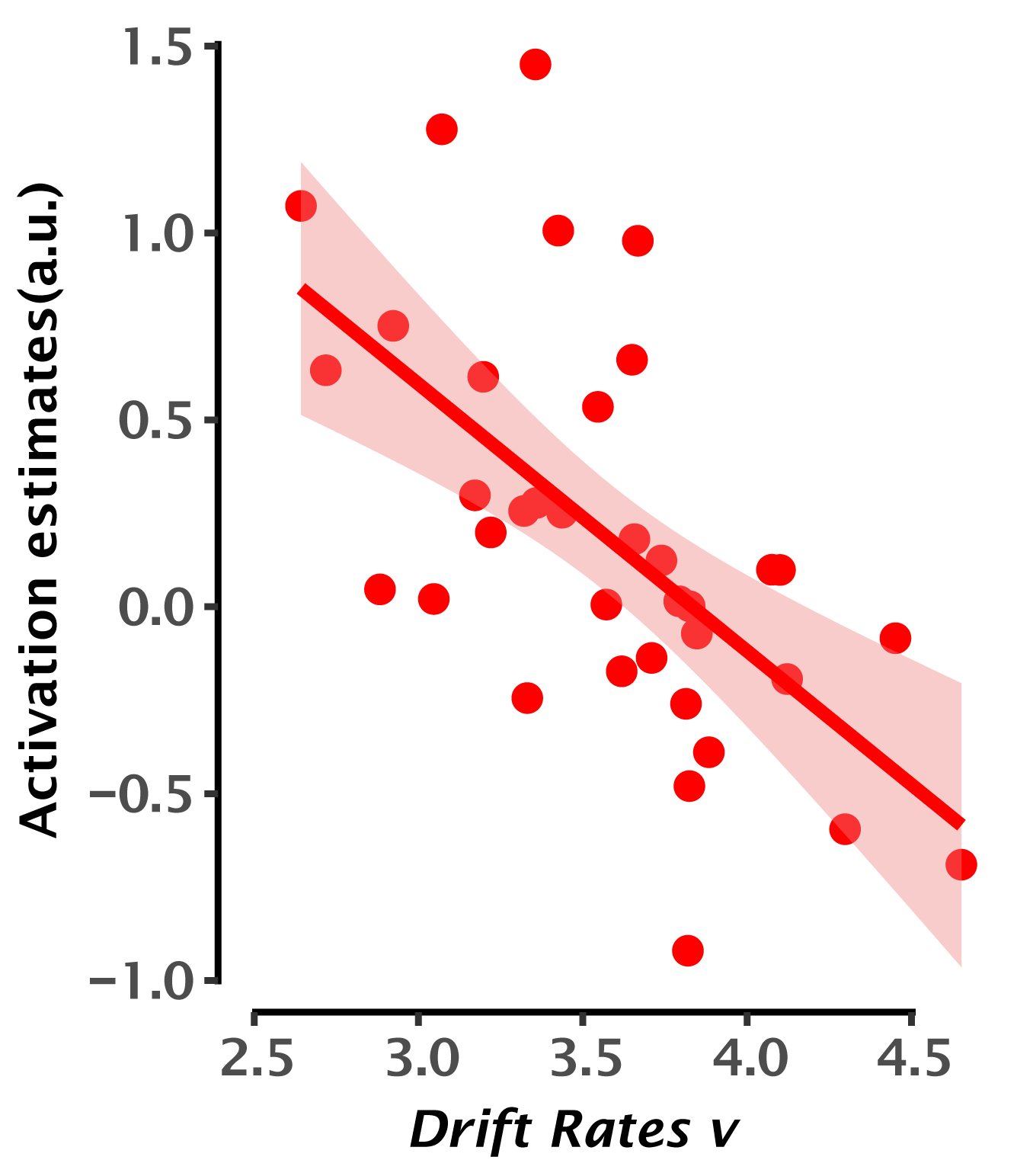

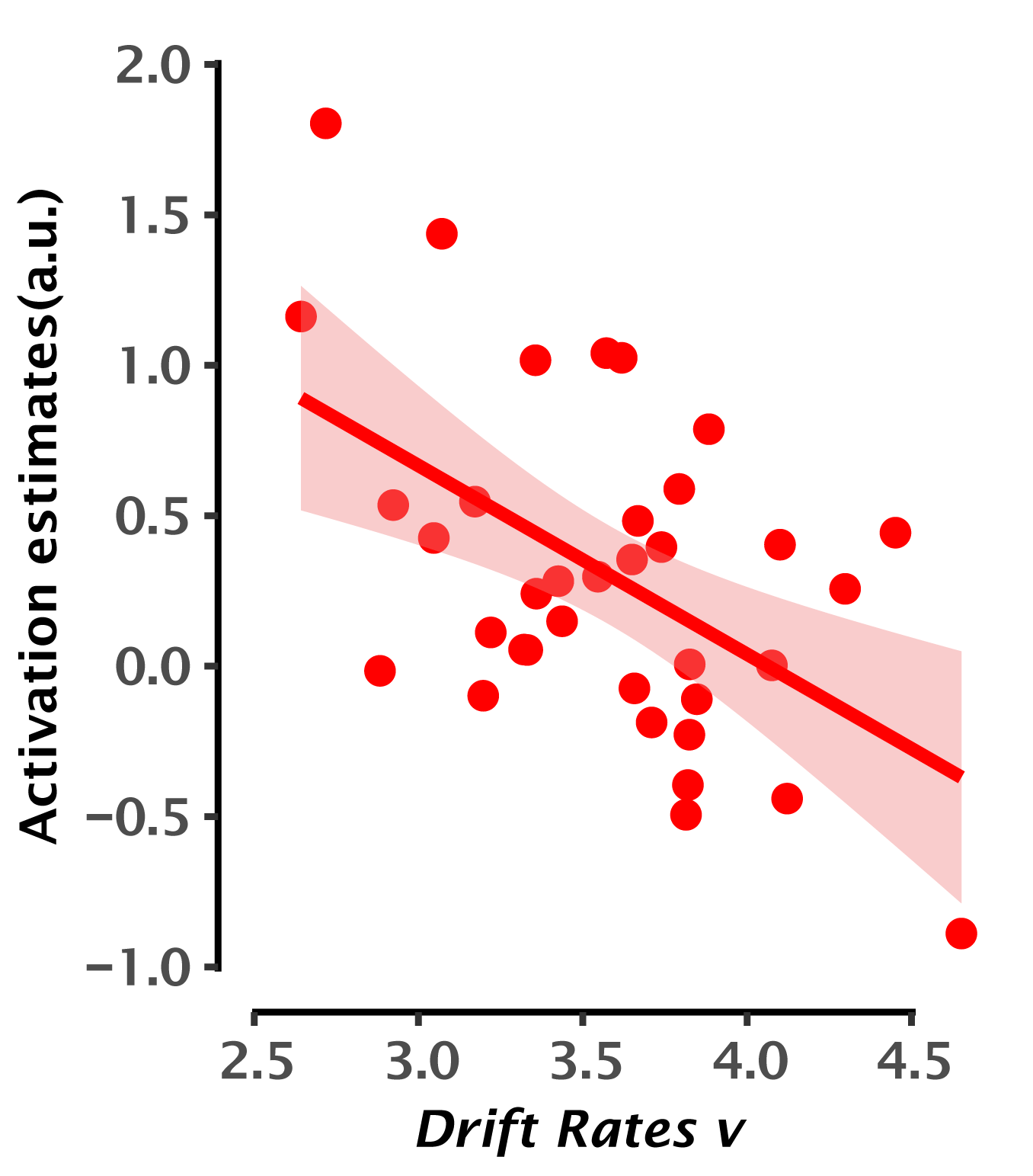

**Figure S10. Brain regions linked with individual differences in drift rate under long-term stress.** Upper panels: Representative slices of significant clusters in the right intraparietal sulcus (IPS), the left superior frontal gyrus (SFG) and the right middle frontal gyrus (MFG). These clusters were derived a whole-brain regression analysis for the contrast map of 2- vs. 0-back condition drift rate as a covariate of interest, with a height threshold *p* < 0.005, an extent threshold *p* < 0.05 corrected. Lower panels: Scatter plots depict the correlations of WM-related activity in the IPS, SFG and MFG with the drift rate under long-term stress. Color bar indicates minimum and maximal T values.

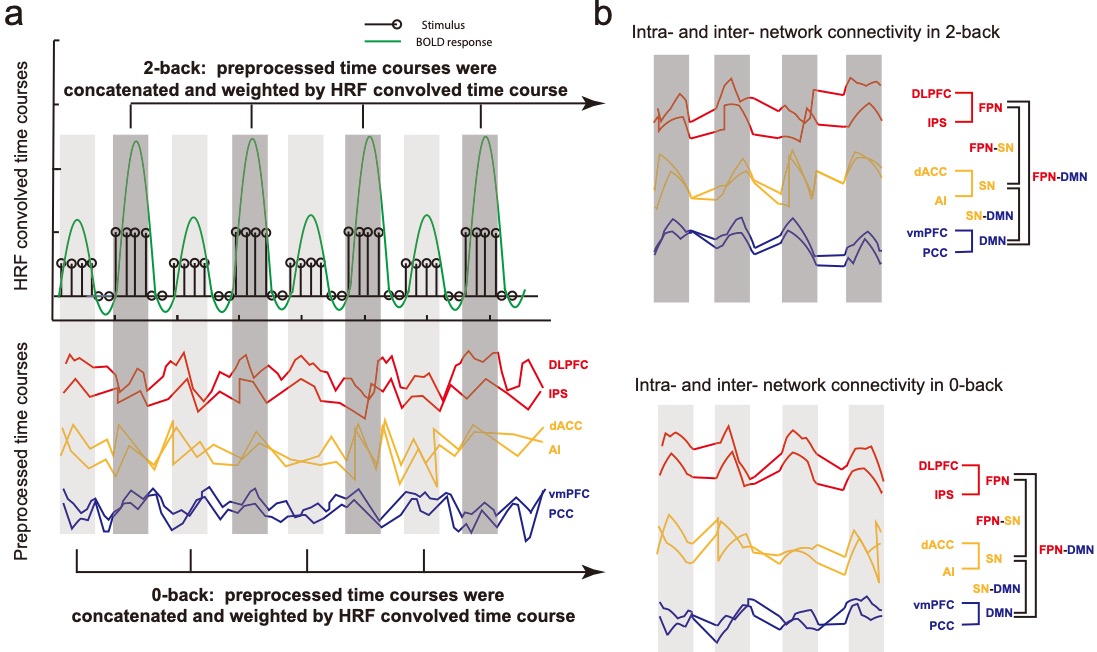

**Figure S11. An illustration of calculating intra- and inter-network connectivity.** (**a**) Preprocessed time courses of six regions of interest (ROIs) and hemodynamic response function (HRF) convolved time course in working memory task. (**b**) Concatenated and weighted time course used to calculate intra- and inter-network functional connectivity strength. Notes: FPN: frontal-parietal network; SN: salience network; DMN: default-node network; dlPFC, dorsal-lateral prefrontal cortex; IPS, intra-parietal sulcus; dACC, dorsal anterior cingulate cortex; vmPFC, ventromedial prefrontal cortex; PCC, posterior cingulate cortex.

**Supplementary Tables: S1 to S22**

**Table S1. Descriptive statistics of trait anxiety and psychological distress (SCL) in stress and control groups**

|  |  | ***Stress*** | ***Control*** |
| --- | --- | --- | --- |
|  | N | 36 | 32 |
| Trait anxiety | Mean ± S.D. | 39.22 ± 7.95 | 41.84 ± 8.36 |
|  | Range | 21 - 52 | 22 - 58 |
|  | Skew | -0.38 | -0.27 |
|  | Kurtosis | -0.78 | -0.58 |
| SCL | Mean ± S.D. | 5.37 ± 3.05 | 4.11 ± 3.59 |
|  | Range | 1.14 – 11.67 | 0 – 13.88 |
|  | Skew | 0.49 | -0.76 |
|  | Kurtosis | 1.21 | 0.73 |

Notes: S.D., standard deviation; SCL, Symptom Checklist 90.

**Table S2. Correlations among trait anxiety, psychological distress (SCL) and state anxiety**

| ***Models*** | ***Stress*** | | ***Control*** | | ***Stress vs. Control*** | |
| --- | --- | --- | --- | --- | --- | --- |
|  | r(34) | p | r(29) | p | z | p |
| Triat anxiety ~ state anxiety | 0.71 | < 0.001 | 0.69 | < 0.001 | 0.15 | 0.88 |
| SCL ~ trait anxiety | 0.65 | < 0.001 | 0.41 | 0.02 | 1.32 | 0.19 |
| SCL ~ state anxiety | 0.50 | 0.002 | 0.10 | 0.59 | 1.75 | 0.08 |
| SCL ~ trait anxiety  (partial out state anxiety) | 0.48 | 0.004 | 0.47 | 0.008 | 0.05 | 0.96 |

Notes: SCL, Symptom Checklist 90. One outlier larger than 2.5 standard deviations for the SCL data was excluded from these correlations in the control group.

**Table S3. Gelman-Rubin R hat statistics of decision parameters**

| ***Parameters*** | ***Stress*** | | ***Control*** | |
| --- | --- | --- | --- | --- |
|  | 0-back | 2-back | 0-back | 2-back |
| v | 0.9999495 | 1.0000673 | 1.0034398 | 1.0038763 |
| a | 1.0001556 | 1.0001203 | 1.0058023 | 1.0109975 |
| t | 0.9999470 | 1.0000809 | 1.0041615 | 1.0087788 |
| z | 1.000006 ~ 1.004565 | | 0.9999727 ~ 1.0035089 | |

Notes: v, drift rate; a, boundary or decision threshold; t, non-decision time; z, start point bias.

**Table S4. Statistics for the effects of long-term stress on all of latent parameters**

|  | ***v*** | | ***a*** | | ***t*** | | ***z*** | |
| --- | --- | --- | --- | --- | --- | --- | --- | --- |
|  | F | P | F | P | F | P | t | P |
| Group | 8.91 | 0.004 | 4.15 | 0.046 | 0.26 | 0.62 | - | - |
| WM load | 629.17 | <0.001 | 36.75 | <0.001 | 131.52 | <0.001 | - | - |
| Group*WM | 0.028 | 0.87 | 1.93 | 0.17 | 0.08 | 0.78 | - | - |
| Stress vs. Control | - | | | | | | 0.086 | 0.93 |

Notes: The main effects of Group and WM loads as well as their interaction effects for latent computational measures (i.e., drift rate *v*, decision threshold *a*, non-decision time *t*), by conducting 2 (Group: stress vs. control)-by-2 (WM load: 0- vs. 2-back) ANOVAs. Each participant’s response bias *z* is assumed to not vary as a function of within-subject conditions (i.e., 0- and 2-back here). Two sample t-test was only conducted for the response bias *z* to examine the group difference. Dashed line indicates not applicable.

**Table S5. Correlations among behavioral measures, trait anxiety and state anxiety**

| ***Models*** | ***Stress*** | | | | | | ***Control*** | | | | | | |
| --- | --- | --- | --- | --- | --- | --- | --- | --- | --- | --- | --- | --- | --- |
|  | ***0-back 2-back*** | | | | | | ***0-back 2-back*** | | | | | | |
|  | r(34) | p | r(34) | p | r(30) | | | p | | | r(30) | | p |
| RT ~ trait anxiety | -0.05 | 0.80 | -0.08 | 0.63 | 0.21 | | | 0.25 | | | 0.14 | | 0.45 |
| RT ~ state anxiety | -0.05 | 0.75 | -0.16 | 0.36 | 0.12 | | | 0.52 | | | 0.05 | | 0.78 |
| RT ~ trait anxiety  (partial out state anxiety) | -0.01 | 0.95 | 0.04 | 0.82 | 0.18 | | | 0.33 | | | 0.14 | | 0.44 |
| Accuracy ~ trait anxiety | 0.17 | 0.33 | 0.05 | 0.79 | -0.28 | | | 0.12 | | | -0.30 | | 0.09 |
| Accuracy ~ state anxiety | 0.13 | 0.45 | 0.17 | 0.31 | -0.38 | | | 0.03 | | | -0.29 | | 0.11 |
| Accuracy ~ trait anxiety  (partial out state anxiety) | 0.11 | 0.54 | -0.11 | 0.52 | -0.03 | | | 0.87 | | | -0.15 | | 0.41 |
| v ~ trait anxiety | -0.30 | 0.08 | -0.09 | 0.60 | -0.08 | | | 0.68 | | | -0.05 | | 0.80 |
| v ~ state anxiety | -0.24 | 0.17 | -0.01 | 0.96 | 0.06 | | | 0.75 | | | -0.15 | | 0.40 |
| v ~ trait anxiety  (partial out sate anxiety) | -0.18 | 0.29 | -0.12 | 0.49 | -0.16 | | | 0.38 | | | 0.09 | | 0.65 |
| a ~ trait anxiety | -0.06 | 0.74 | -0.10 | 0.56 | | 0.15 | | | 0.40 | 0.06 | | 0.76 | |
| a ~ state anxiety | -0.09 | 0.59 | -0.14 | 0.43 | | 0.001 | | | 1.00 | -0.11 | | 0.56 | |
| a ~ trait anxiety  (partial out state anxiety) | 0.01 | 0.95 | -0.006 | 0.97 | | 0.21 | | | 0.25 | 0.18 | | 0.33 | |

Notes: RT, reaction time; state, state anxiety; v, drift rate; a, decision threshold; r, Pearson’s correlation coefficients

**Table S6. Parameters and minimum cluster size from Monte-Carlo simulations**

|  | **P(uncorrected)** | **voxels** | **Minimun voxels needed to surpass p< 0.05** | **Mask** | **P(corrected)** |
| --- | --- | --- | --- | --- | --- |
| Insula | 0.001 | 240 | 34 | Whole brain gray matter | < 0.05 |
| IPS ^a^ | 0.005 | 85 | 65 |  | < 0.05 |
| PCC ^a^ | 0.005 | 215 | 65 |  | < 0.05 |
| AI ^a^ | 0.005 | 67 | 65 | Whole brain gray matter | < 0.05 |
| MFG ^a^ | 0.005 | 56 | 45 | PFC mask | < 0.05 |
| mPFC ^a^ | 0.005 | 46 | 45 |  | < 0.05 |
| dACC ^a^ | 0.005 | 45 | 30 | Cingulate mask | < 0.05 |

Notes: The initial height threshold for our fMRI analyses was set at *p < 0.001* on the voxel-wise level and an extent threshold *p* < 0.05 on the cluster-wise level corrected for multiple comparisons using suprathreshold cluster-size approach based on Monte-Carlo simulations.

^a^ Given our priori hypotheses regarding the DMN, SN and FPN, these regions were additionally investigated using a height threshold of *p < 0.005* voxel-vise and an extent threshold of *p < 0.05* cluster-wise corrected for multiple comparisons.

**Table S7. Statistics of the mediation models for trait anxiety, IPS/MFG and drift rate**

| ***Models*** | ***Stress*** | | ***Control*** | | | ***Stress vs. Control*** | |
| --- | --- | --- | --- | --- | --- | --- | --- |
|  | β | 95% CI | β | 95% CI | | β | 95% CI |
| Mediation model: trait anxiety - IPS - drift rate | | | | | | | |
| IPS ~ trait anxiety | 0.035 | [0.017, 0.052] | 0.007 | [-0.018, 0.025] |  | |  |
| v ~ IPS | -0.454 | [-0.794, -0.237] | 0.358 | [0.079, 0.684] |  | |  |
| v ~ trait anxiety | 0.011 | [-0.008, 0.031] | -0.005 | [-0.023,0.014] |  | |  |
| Indirect Est. | -0.016 | [-0.034, -0.006] | 0.002 | [-0.004, 0.014] | -0.018 | | [-0.038, -0.005] |
| Mediation model: trait anxiety - MFG - drift rate | | | | | | | |
| MFG ~ trait anxiety | 0.027 | [0.01, 0.047] | -0.011 | [-0.04,0.011] |  | |  |
| v ~ MFG | -0.474 | [-0.685, -0.199] | 0.124 | [-0.092, 0.292] |  | |  |
| v ~ trait anxiety | 0.008 | [-0.009, 0.024] | -0.001 | [-0.023, 0.018] |  | |  |
| Indirect Est. | -0.013 | [-0.029, -0.003] | -0.001 | [-0.008, 0.001] | -0.012 | | [-0.028, -0.001] |

Notes: β, regression weight; CI, confidence interval; Est., estimation. MFG, middle frontal gyrus; IPS, intra-parietal sulcus; v, drift rate. The indirect effect was considered significant if the 95% CI did not include zero.

**Table S8. Statistics for the mediation models of state anxiety, IPS/MFG and drift rate**

| ***Models*** | ***Stress*** | | ***Control*** | | ***Stress vs. Control*** | |
| --- | --- | --- | --- | --- | --- | --- |
|  | β | 95% CI | β | 95% CI | β | 95% CI |
| Mediation model: state anxiety - IPS - drift rate | | | | | | |
| IPS ~ state anxiety | 0.028 | [0.014, 0.048] | -0.017 | [-0.038, 0.004] |  |  |
| v ~ IPS | -0.461 | [-0.764, -0.243] | 0.338 | [0.02, 0.644] |  |  |
| v ~ state anxiety | 0.011 | [-0.013, 0.029] | -0.002 | [-0.023, 0.016] |  |  |
| Indirect Est. | -0.013 | [-0.026, -0.006] | -0.006 | [-0.018, 0] | -0.007 | [-0.020, 0.006] |
| Mediation model: state anxiety - MFG - drift rate | | | | | | |
| MFG ~ state anxiety | 0.022 | [0.003, 0.048] | -0.022 | [-0.045, 0.001] |  |  |
| v ~ MFG | -0.482 | [-0.717, -0.225] | 0.105 | [-0.143, 0.283] |  |  |
| v ~ state anxiety | 0.009 | [-0.012, 0.022] | -0.006 | [-0.027, 0.015] |  |  |
| Indirect Est. | -0.01 | [-0.028, -0.001] | -0.002 | [-0.011, 0.002] | -0.008 | [-0.025, 0.003] |

Notes: β, regression weight; CI, confidence interval; Est., estimation. MFG, middle frontal gyrus; IPS, intra-parietal sulcus; v, drift rate. The indirect effect was considered significant if the 95% CI did not include zero.

**Table S9. Relations of latent-, neural-, and connectivity-measures with WM accuracy**

| ***Models*** | ***Stress Control*** | | | |
| --- | --- | --- | --- | --- |
|  | r(34) | p | r(30) | p |
| Accuracy ~ v | 0.24 | 0.15 | 0.27 | 0.14 |
| Accuracy ~ a | 0.28 | 0.09 | -0.08 | 0.64 |
| Accuracy ~ Insula_L | -0.32 | 0.06 | 0.33 | 0.07 |
| Accuracy ~ Insula_R | -0.48 | 0.003 | 0.15 | 0.42 |
| Accuracy ~ SN_DMN | -0.49 | 0.002 | 0.15 | 0.42 |
| Accuracy ~ FPN_DMN | 0.20 | 0.25 | -0.20 | 0.27 |
| Accuracy ~ PCC | 0.29 | 0.09 | 0.01 | 0.95 |
| Accuracy ~ mPFC | -0.18 | 0.29 | 0.36 | 0.04 |
| Accuracy ~ insula | -0.27 | 0.12 | 0.33 | 0.06 |
| Accuracy ~ dACC | -0.10 | 0.55 | 0.17 | 0.36 |
| Accuracy ~ IPS | -0.10 | 0.57 | 0.23 | 0.21 |
| Accuracy ~ MFG | -0.37 | 0.03 | 0.06 | 0.74 |

Notes: The correlation analyses were mainly restricted on behavioral, latent, neural and connectivity metrics in the 2-back condition.

**Table S10. The relationship of latent-, neural-, and connectivity-measures with RTs**

| ***Models*** | ***Stress*** | | | ***Control*** |
| --- | --- | --- | --- | --- |
|  | r(34) | p | r(30) | p |
| RT ~ v | -0.43 | 0.008 | -0.68 | <0.001 |
| RT ~ a | 0.94 | <0.001 | 0.81 | <0.001 |
| RT ~ Insula_L | -0.01 | 0.93 | -0.19 | 0.30 |
| RT ~ Insula_R | -0.13 | 0.44 | 0.23 | 0.21 |
| RT ~ SN_DMN | -0.30 | 0.08 | -0.05 | 0.78 |
| RT ~ FPN_DMN | -0.12 | 0.49 | 0.03 | 0.86 |
| RT ~ PCC | -0.14 | 0.41 | -0.09 | 0.64 |
| RT ~ mPFC | -0.18 | 0.30 | -0.12 | 0.53 |
| RT ~ insula | 0.11 | 0.52 | -0.28 | 0.12 |
| RT ~ dACC | -0.20 | 0.24 | -0.06 | 0.73 |
| RT ~ IPS | 0.05 | 0.76 | -0.16 | 0.40 |
| RT ~ MFG | -0.02 | 0.89 | -0.21 | 0.26 |

Notes: These correlation analyses were mainly restricted on behavioral, latent, neural and connectivity metrics in the 2-back condition. SN, salience network; DMN, default mode network; PCC, posterior cingulate cortex; mPFC, medial prefrontal cortex; dACC, dorsal anterior cingulate cortex; IPS, intraparietal sulcus; MFG, middle frontal gyrus.

**Table S11. Summary outcomes from the prediction analyses**

| ***Models*** | ***Stress*** | | ***Control*** | |
| --- | --- | --- | --- | --- |
|  | r_(predicted, observed)_ | p | r_(predicted, observed)_ | p |
| SCL ~ trait anxiety | 0.61 | < 0.001 | 0.35 | 0.010 |
| SCL ~ trait anxiety  (controlling state anxiety) | 0.46 | 0.002 | 0.31 | 0.022 |
| RT ~ v | 0.48 | < 0.001 | 0.67 | < 0.001 |
| RT ~ a | 0.91 | < 0.001 | 0.80 | < 0.001 |
| IPS ~ trait anxiety | 0.33 | < 0.001 | - | - |
| MFG ~ trait anxiety | 0.27 | 0.016 | - | - |
| v ~ IPS | 0.36 | 0.002 | 0.28 | 0.014 |
| v ~ MFG | 0.42 | 0.002 | - | - |
| FPN_DMN ~ triat anxiety  (controlling state anxiety) | 0.13 | 0.10 | - | - |

Notes: Dashes represent that prediction analyses were not performed, because non-significant correlations were observed from conventional correlational analyses. SCL, psychological distress measure by Symptom Checklist; v, drift rate; a, decision threshold; IPS, intraparietal sulcus; MFG, middle frontal gyrus; FPN, frontoparietal network; DMN, default mode network.

**Table S12. Brain regions showing the main effect of long-term stress**

| Region | MNI coordinates | | | T value | Cluster size |
| --- | --- | --- | --- | --- | --- |
|  | x | y | z |  |  |
| *Long-term stress > Control group* | | | | | |
| Anterior insula L | -26 | 14 | 16 | 4.49 | 240 |
| Anterior insula R | 38 | 20 | 14 | 3.93 | 156^a^ |
| PCC R | 12 | -32 | 28 | 3.64 | 114^a^ |
| Middle occipital cortex extending into the cuneus | 6 | -74 | 30 | 5.45 | 1904 |
| Temporoparietal junction L | -48 | -46 | 24 | 4.07 | 54 |
| *Control group > Long-term stress* | | | | | |
| Orbital PFC L | -34 | 38 | -10 | 3.88 | 40 |

Notes: Significant clusters were determined by using a height threshold of p < 0.001 and an extent threshold of p < 0.05 corrected according to suprathreshold cluster-size approach based on Monte-Carlo simulations. ^a^ Given our a priori hypotheses on regions of interest in the FPN, DMN and SN, we applied a height threshold of p < 0.005 and a spatial extent threshold of p < 0.05 corrected for these regions. PCC, posterior cignulate cortex; L, left; R, right.

**Table S13. Brain WM-related activation/deactivation showing interaction between Group (long-term stress vs. control) and WM-load (2-back vs. 0-back)**

| Region | MNI coordinates | | | T value | Cluster size |
| --- | --- | --- | --- | --- | --- |
|  | x | y | z |  |  |
| *WM-related activation (2- > 0-back) vs. (Long-term stress > Control)* | | | | | |
| Anterior insula L | -34 | 10 | 12 | 3.28 | 58^a^ |
| dACC R | 8 | 18 | 28 | 3.25 | 45^a^ |
| *WM-related deactivation (0- > 2-back) vs. (Long-term stress > Control)* | | | | | |
| ACC | 4 | 34 | 10 | 3.44 | 100^a^ |
| PCC | 10 | -46 | 26 | 3.37 | 215^a^ |
| PCC | -6 | -30 | 30 | 3.51 | 80^a^ |
| Medial prefrontal cortex | 16 | 54 | 4 | 4.12 | 46^a^ |
| Posterior insula/Superior temporal gyrus R | 56 | -2 | 0 | 4.42 | 1142 |
| Paracentral lobe L | -4 | -38 | 54 | 3.72 | 72 |
| Temporoparietal junction L | -52 | -22 | 16 | 3.50 | 268 |
| Superior occipital lobe R | 24 | -82 | 30 | 4.36 | 247 |

Notes: Significant clusters were determined by using a height threshold of p < 0.001 and an extent threshold of p < 0.05 corrected according to suprathreshold cluster-size approach based on Monte-Carlo simulations. ^a^ Given our a priori hypotheses on regions of interest in the FPN, DMN and SN, we applied a height threshold of p < 0.005 and a spatial extent threshold of p < 0.05 corrected for these regions. PCC, posterior cignulate cortex; ACC, anterior cingulate cortex; dACC, dorsal anterior cingulate cortex; mPFC, medial prefrontal cortex; L, left; R, right.

**Table S14. The correlations between activation in the insula and drift rate**

| ***Models*** | ***Stress*** | | | | | | ***Control*** | | |
| --- | --- | --- | --- | --- | --- | --- | --- | --- | --- |
|  | ***0 back 2 back*** | | | | ***0 back 2 back*** | | | | |
|  | r(34) | p | r(34) | p | r(30) | p | | r(30) | p |
| Insula_L ~ v | 0.02 | 0.90 | 0.07 | 0.69 | -0.14 | 0.44 | | 0.05 | 0.78 |
| Insula_R ~ v | 0.005 | 0.98 | -0.13 | 0.45 | -0.13 | 0.48 | | -0.12 | 0.50 |

**Table S15. Brain regions linked to drift rate in 0-, 2- and 2- vs. 0-back contrasts in long-term stress group**

| Regions | MNI coordinates | | | T value | Cluster size |
| --- | --- | --- | --- | --- | --- |
|  | x | y | z |  |  |
| *0-back* | | | | | |
| Superior Frontal lobe L | -24 | 22 | 56 | 3.60 | 56 ^a^ |
| *2-back* | | | | | |
| Superior Frontal lobe L | -24 | 56 | 12 | 4.84 | 243 ^a^ |
| IPS R | 32 | -48 | 44 | 3.54 | 80 ^a^ |
| MFG R | 46 | 36 | 22 | 2.98 | 56 ^a^ |
| *2- versus 0-back* | | | | | |
| No brain regions were observed. | | | | | |

Notes: ^a^ Given our a priori hypotheses on regions of interest in the FPN, DMN and SN regions, we applied a height threshold of p < 0.005 and a spatial extent threshold of p < 0.05 corrected for these regions. IPS, intraparietal sulcus; MFG, middle frontal gyrus.

**Table S16. Brain regions linked to drift rate in 0-, 2- and 2- vs. 0-back** **conditions in the control group**

| Regions | MNI coordinates | | | T value | Cluster size |
| --- | --- | --- | --- | --- | --- |
|  | x | y | z |  |  |
| *0-back* | | | | | |
| Superior Frontal lobe R | 16 | 44 | 26 | 3.69 | 97 ^a^ |
| *2-back* | | | | | |
| Superior Frontal lobe R | 26 | -62 | 42 | 4.51 | 75 ^a^ |
| *2- versus 0-back* | | | | | |
| Frontal lobe / White matter L | -30 | 38 | 0 | 4.09 | 154 ^a^ |
| Frontal lobe / White matter R | 22 | 42 | -2 | 4.04 | 118 ^a^ |

Notes: ^a^ Given our a priori hypotheses on regions of interest in the FPN, DMN and SN regions, we applied a height threshold of p < 0.005 and a spatial extent threshold of p < 0.05 corrected for these regions.

**Table S17. Correlations between neural activity in the SN regions and psychological distress**

| ***Models*** | ***Stress*** | | | | ***Control*** | | | |
| --- | --- | --- | --- | --- | --- | --- | --- | --- |
|  | ***0 back 2 back*** | | | | ***0 back 2 back*** | | | |
|  | r(34) | p | r(34) | p | r(29) | p | r(29) | p |
| *Long-term stress vs. Control* | | | | | | | | |
| Insula_L ~ SCL | 0.16 | 0.36 | 0.30 | 0.072 | -0.09 | 0.61 | -0.06 | 0.75 |
| Insula_R ~ SCL | 0.23 | 0.17 | 0.29 | 0.088 | -0.25 | 0.17 | -0.05 | 0.79 |
| *WM-related activation (2- vs. 0-back) vs. (Long-term stress vs. Control)* | | | | | | | | |
| Insula_L ~ SCL | 0.21 | 0.22 | 0.23 | 0.18 | -0.05 | 0.79 | -0.10 | 0.59 |
| dACC ~ SCL | -0.02 | 0.89 | -0.30 | 0.08 | -0.01 | 0.96 | 0.11 | 0.54 |

**Table S18. Brain regions linked to psychological distress in 0-, 2- and 2- vs. 0-back conditions in the long-term stress group**

| Regions | MNI coordinates | | | T values | Cluster size |
| --- | --- | --- | --- | --- | --- |
|  | x | y | z |  |  |
| *0-back* | | | | | |
| No brain region was found | | | | | |
| *2-back* | | | | | |
| Posterior cingulate gyrus | 14 | -28 | 50 | 5.19 | 687 |
| Medial Frontal Gyrus | -2 | 62 | 24 | 5.74 | 443^a^ |
| Inferior frontal gyrus | -36 | 22 | -18 | 5.38 | 91^a^ |
| *2- vs. 0-back* | | | | | |
| Superior medial frontal gyrus | 12 | 56 | 26 | 4.31 | 120 ^a^ |
| Middle frontal gyrus R | 40 | 46 | -10 | 4.87 | 70 ^a^ |
| Middle cingulate gyrus | -4 | -6 | 36 | 3.97 | 61 |
| Corpus callosum | -12 | 30 | 6 | 4.43 | 106 |

Notes: Significant clusters were determined by using a height threshold of p < 0.001 and an extent threshold of p < 0.05 corrected according to suprathreshold cluster-size approach based on Monte-Carlo simulations. ^a^ We applied a height threshold of p < 0.005 and a spatial extent threshold of p < 0.05 corrected only for an explortory purpose.

**Table S19. Brain regions linked with psychological distress in 0-, 2- and 2- vs. 0-back** **conditions in the control group**

| Regions | MNI coordinates | | | T values | Cluster size |
| --- | --- | --- | --- | --- | --- |
|  | x | y | z |  |  |
| *0-back* | | | | | |
| PCC R | 12 | -56 | 10 | 4.64 | 111 ^a^ |
| *2-back* | | | | | |
| PCC L | -8 | -54 | 6 | 4.18 | 103 ^a^ |
| PCC R | 10 | -56 | 8 | 4.68 | 210 ^a^ |
| Middle frontal gyrus R | 44 | 50 | -12 | 5.09 | 226 ^a^ |
| Inferior parietal lobe L | -44 | -58 | 44 | 3.78 | 77 ^a^ |
| Inferior temporal gyrus R | 58 | -18 | -24 | 5.19 | 65 |
| Superior temopral gyrus R | 50 | -54 | 22 | 4.62 | 68 |
| *2- vs. 0-back* | | | | | |
| Middle frontal gyrus R | 46 | 50 | -10 | 4.34 | 227 ^a^ |
| Middle frontal gyrus L | -28 | 48 | -8 | 4.01 | 81 ^a^ |
| Middle frontal gyrus L | -50 | 42 | -2 | 4.22 | 80 ^a^ |
| Angular L | -50 | -58 | 32 | 3.45 | 64 ^a^ |
| Middle cingulate gyrus | 2 | -26 | 38 | 5.20 | 46 |
| Middle Temporal gyrus R | 58 | -26 | -14 | 4.32 | 38 |

Notes: Significant clusters were determined by using a height threshold of p < 0.001 and an extent threshold of p < 0.05 corrected according to suprathreshold cluster-size approach based on Monte-Carlo simulations. ^a^ We applied a height threshold of p < 0.005 and a spatial extent threshold of p < 0.05 corrected only for an exploratory purpose.

**Table S20. MNI coordinates of the FPN, SN and DMN nodes for the network analysis**

| ***Brain regions*** | ***x*** | ***y*** | ***z*** |
| --- | --- | --- | --- |
| PCC ^a^ | -4 | -52 | 22 |
| mPFC ^a^ | -2 | 50 | 18 |
| vACC/vmPFC ^a^ | 2 | 32 | -8 |
| DLPFC L ^b^ | -40 | 26 | 24 |
| DLPFC R ^b^ | 40 | 32 | 30 |
| IPS L ^b^ | -36 | -50 | 40 |
| IPS R ^b^ | 40 | -48 | 38 |
| dACC ^c^ | -6 | 18 | 30 |
| Insula L ^c^ | -40 | 18 | -12 |
| Insula R ^c^ | 42 | 10 | -12 |

Notes: ^a^ Nodes in the DMN. ^b^ Nodes in the FPN. ^c^ Nodes in the SN. PCC, posterior cignulate cortex; mPFC, medial prefrontal cortex; vACC, ventral anterior cingulate cortex; DLPFC, dorsolateral prefrontal cortex; IPS, intraparietal sulcus; dACC, dorsal anterior cingulate cortex; L, left; R, right

**Table S21. Effects of both intra- and inter connectivity**

|  | ***FPN*** | | ***DMN*** | | ***SN*** | | ***FPN-DMN*** | | ***FPN-SN*** | | ***SN-DMN*** | |
| --- | --- | --- | --- | --- | --- | --- | --- | --- | --- | --- | --- | --- |
|  | F | P | F | P | F | P | F | P | F | P | F | P |
| Group | 1.70 | 0.20 | 0.78 | 0.38 | 1.69 | 0.20 | 5.04 | 0.028 | 0.59 | 0.45 | 2.55 | 0.12 |
| Condition | 4.17 | 0.05 | 2.47 | 0.12 | 0.82 | 0.37 | 12.79 | 0.001 | 1.41 | 0.24 | 4.67 | 0.03 |
| Interaction | 1.11 | 0.30 | 3.07 | 0.08 | 0.008 | 0.93 | 0.06 | 0.81 | 0.53 | 0.47 | 0.03 | 0.86 |

Notes: Separate 2 (Group: stress vs. control)-by-2 (WM load: 0- vs. 2-back) ANOVAs were conducted for intra- and inter-network coupling metrics among the FPN, DMN and SN regions.

**Table S22. DIC values of 15 model variants**

| ***15 Model variants*** | ***DIC*** | ***DIC relative difference*** |
| --- | --- | --- |
| *z* | -4185 | 532 |
| *t* | -4192 | 525 |
| *tz* | -4305 | 412 |
| *a* | -4383 | 334 |
| *az* | -4465 | 252 |
| *at* | -4575 | 142 |
| *atz* | -4633 | 84 |
| *v* | -4634 | 83 |
| *av* | -4641 | 76 |
| *avt* | -4667 | 50 |
| *vt* | -4674 | 43 |
| *vz* | -4674 | 43 |
| *avz* | -4701 | 16 |
| *vtz* | -4716 | 1 |
| *avtz* | -4717 | 0 |

Notes: *a*, decision threshold; *t*, non-decision time; *v,* drift rate; *z*, start bias.

**References**

1.	Ratcliff R, Rouder JN. Modeling response times for two-choice decisions. *Psychological science* **9**, 347-356 (1998).

2.	Navarro DJ, Fuss IG. Fast and accurate calculations for first-passage times in Wiener diffusion models. *J Math Psychol* **53**, 222-230 (2009).

3.	Matzke D, Wagenmakers E-J. Psychological interpretation of the ex-Gaussian and shifted Wald parameters: A diffusion model analysis. *Psychon B Rev* **16**, 798-817 (2009).

4.	Herz DM, Zavala BA, Bogacz R, Brown P. Neural correlates of decision thresholds in the human subthalamic nucleus. *Curr Biol* **26**, 916-920 (2016).

5.	O’Callaghan C*, et al.* Visual hallucinations are characterized by impaired sensory evidence accumulation: insights from hierarchical drift diffusion modeling in Parkinson’s disease. *Biological Psychiatry: Cognitive Neuroscience and Neuroimaging* **2**, 680-688 (2017).

6.	Herz DM*, et al.* Distinct mechanisms mediate speed-accuracy adjustments in cortico-subthalamic networks. *Elife* **6**, e21481 (2017).

7.	Owen AM, McMillan KM, Laird AR, Bullmore E. N-back working memory paradigm: a meta-analysis of normative functional neuroimaging studies. *Hum Brain Mapp* **25**, 46-59 (2005).

8.	Laird AR, Eickhoff SB, Li K, Robin DA, Glahn DC, Fox PT. Investigating the functional heterogeneity of the default mode network using coordinate-based meta-analytic modeling. *J Neurosci* **29**, 14496-14505 (2009).

9.	Seeley WW*, et al.* Dissociable Intrinsic Connectivity Networks for Salience Processing and Executive Control. *The Journal of Neuroscience* **27**, 2349 (2007).
